# Epistasis and the changing fitness landscapes of SARS-CoV-2

**DOI:** 10.64898/2026.03.12.711354

**Authors:** Luca Sesta, Richard A. Neher

## Abstract

Since its emergence in late 2019, millions of SARS-CoV-2 genomes have been generated as part of global efforts to monitor the evolution and spread of the virus. This unprecedented volume of data provides a unique opportunity to study viral evolution at unparalleled resolution. In particular, individual genomic sites can be observed to have mutated independently thousands of times. These mutation counts have been used to estimate site-specific mutation rates and fitness effects for most mutations across the viral genome. Here, we use these data to investigate how the landscape of mutational fitness costs has changed over the course of the pandemic. SARS-CoV-2 evolution over the past six years has been characterized by the emergence of distinct variants separated by long branches corresponding to evolutionary saltations involving up to 50 mutations. We compare inferred fitness landscapes of the Spike protein across these variants and find that shifts in the estimated effects of non-synonymous mutations are linked to genetic differences between them. Sites with altered fitness costs are enriched near positions where the genetic backgrounds differ. To explain the observed changes, we introduce a model with pairwise epistatic interactions between mutations and residues that differ between variants. This model is able to explain about half of the variance in the shifts of fitness effects and suggests that each mismatch between variants substantially alters mutation effects at typically 1 to 3 additional positions.

## I. INTRODUCTION

The Severe Acute Respiratory Syndrome Coronavirus 2 (SARS-CoV-2) is a positive sense RNA virus which entered the human population in 2019 and caused the global COVID-19 pandemic in the years that followed. While vaccination reduced morbidity and mortality associated with the virus in regions where it was available early enough, continued evolution of the virus led to the emergence of multiple variants that were able to partially evade pre-existing immunity. This rapid evolution motivated global genomic surveillance efforts such that the emergence and spread of novel variants could be followed in near real time at unprecedented resolution. Fig. 1A shows a phylogenetic tree of a few thousand representative viral sequences from the beginning of the pandemic up to November 2024 – less than 0.1% of all available data (Hadfield *et al*., 2018). To this day, there are approximately *∼*18 million full genome sequences available in the GISAID database (Shu and McCauley, 2017). These unprecedented data volumes required novel bioinformatic tools and opened up new ways to analyze and track viral evolution. In particular, pandemic scale phylogenetic reconstruction using UShER (McBroome *et al*., 2021; Turakhia et al., 2021) or genetic diversity dashboard like CoV-Spectrum (Chen *et al*., 2022) played a prominent role in monitoring the spread and emergence of viral variants.

**FIG. 1.**
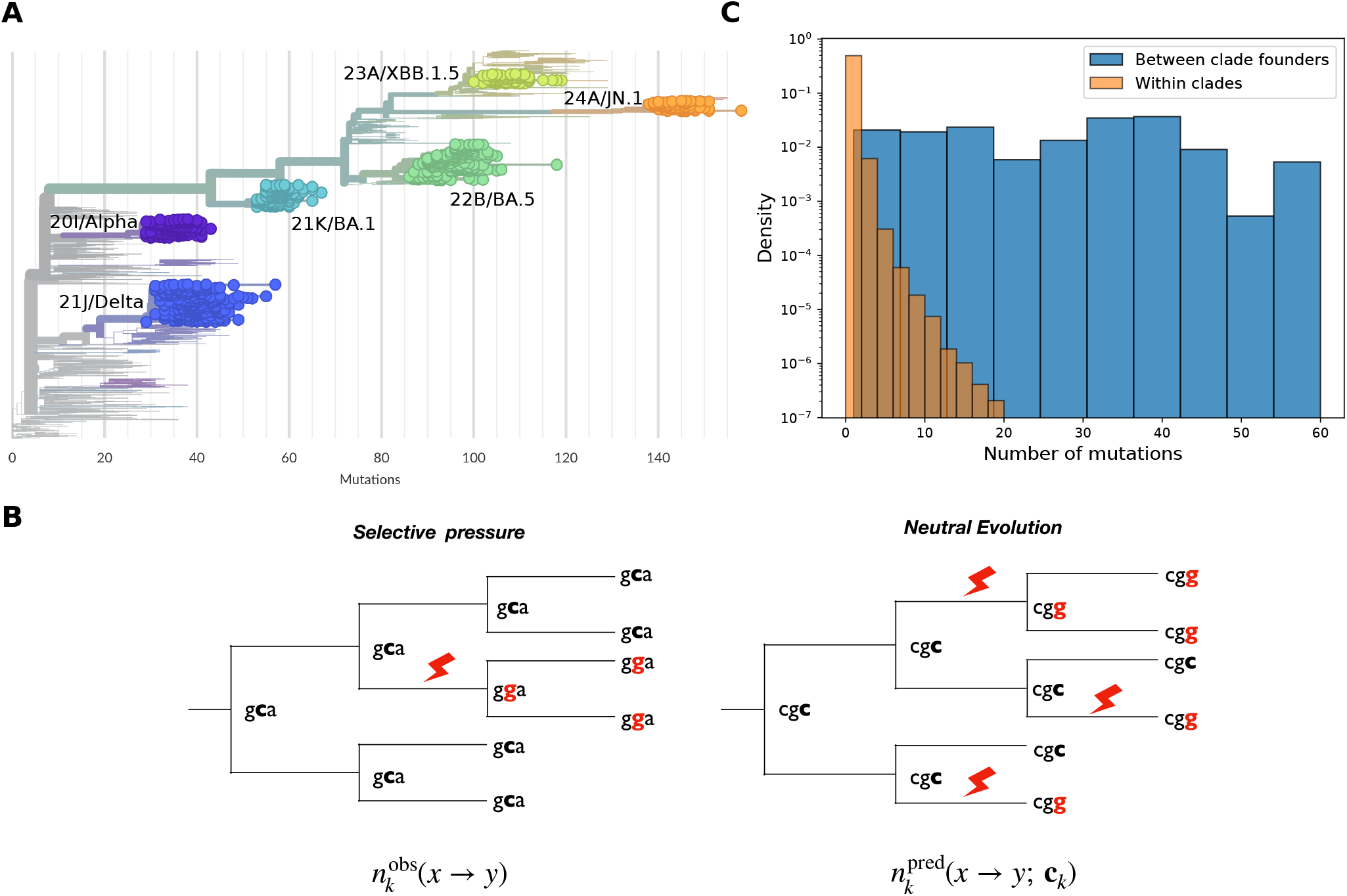
Counting independent mutations on the SARS-CoV-2 phylogeny. **(A)** Phylogenetic tree of the SARS-CoV-2 virus obtained via Nextstrain, up to November 2024. Representative viral clades are labeled and highlighted in distinct colors. The labels include both the Nextstrain and WHO/Pango nomenclature. **(B)** In low diversity samples like SARS-CoV-2 genomes, mutations can be mapped confidently to specific branches of the phylogenetic tree. Independent mutation events can therefore be counted. By comparing observed counts of a mutation (C G in the illustration) at a site under selection (left) to a neutral expectation (right), one can quantify fitness effects of mutations. With a model that predicts mutation counts 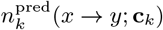 for any mutation type *x → y* in different contexts **c**_*k*_ (Haddox *et al*., 2025b) and observed counts 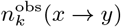, one can estimate the fitness effect Δ*f*_*k*_(*x → y*) of the mutation at site *k*. **(C)** Distance distributions in Spike protein sequences between (blue) and within clades (orange). The former is approximately uniform over the observed range, whereas the latter is concentrated on few mismatches.

At the same time, high-throughput Deep Mutational Scanning (DMS) has been used to systematically assess the impact of mutations on various viral phenotypes such as entry and binding on host-cell surface (Dadonaite *et al*., 2024, 2023; Starr et al., 2020) or immune escape (Cao *et al*., 2023, 2022; Greaney et al., 2021; Starr et al., 2021; Yisimayi *et al*., 2024). Most DMS experiments are restricted to the Receptor Binding Domain (RBD) of the Spike protein which directly interacts with the host receptor of the SARS-CoV-2, the human Angiotensin Converting Enzyme 2 (ACE2), and is the main target of neutralizing antibodies. Notable exceptions are the full-length Spike DMS experiments by Dadonaite et al. (2024, 2023, 2025) and experiments probing the mutational landscapes of the viral protease (Flynn *et al*., 2023; Iketani et al., 2022). The remainder of the viral genome is not readily probed by DMS.

This limitation can be overcome by analyzing patterns of mutations across mutation annotated trees (MATs) of millions of genomes (Turakhia *et al*., 2021). The majority of synonymous mutations are approximately neutral and allow to characterize the mutational spectrum of the virus, i.e. the relative rates of different nucleotide mutations (Bloom *et al*., 2023; De Maio *et al*., 2021; Haddox *et al*., 2025b; Hensel, 2025; Neher, 2022). Bloom and Neher (2023) estimated the fitness effects of most mutations across the genome by comparing the number of independent occurrences of a mutation on the pandemic scale phylogenetic tree to what would be expected from the baseline rates of neutral evolution: the more deleterious a mutation is, the fewer independent occurrences of it will be observed on the tree (see Fig. 1B). Within this framework, sequence data stemming from natural evolution can be used akin to a DMS experiment probing the virus evolution in the wild.

The underlying assumption of these these methods that use mutation counts to estimate fitness effects is that a mutation has the same effect on fitness independently of where it occurs in the tree. Given the low diversity of SARS-CoV-2, this is expected to be approximately true, but changing genetic backgrounds or varying selective pressures exerted by the host immune system could violate this assumption. How fitness landscapes change with the genetic background between different variants of SARS-CoV-2 has been investigated by Starr and collaborators, who compared DMS measurements spanning the RBD in different major viral variants (Starr *et al*., 2022a,b; Taylor and Starr, 2023, 2024). The question of how the fitness landscape changes from one variant to the next is related to but distinct from the role of epistasis between the defining mutations of a variant during its emergence. The latter has been shown to have been critical along the evolutionary trajectory leading to the Omicron variant (Moulana *et al*., 2025, 2022). Specifically, mutations Q498R and N501Y significantly increased binding affinity of Spike to the human ACE2, allowing to buffer the emergence of multiple escape mutations that would have otherwise impaired host-cell binding.

Here, we systematically investigate how the landscape of mutational fitness effects changes in the Spike protein across the SARS-CoV-2 phylogeny. We now have sufficiently many sequences to estimate fitness effects separately in different parts of the phylogenetic tree, i.e. in different clades of closely related variants indicated by different colors in Fig. 1A using the approach by Bloom and Neher (2023). By comparing fitness effect estimates in different clades, we can investigate the phenomenon of *epistasis* and try to explain the observed differences in terms of interactions between mutations and residues that differ between clades. Our approach is based on a minimal epistatic model known as generalized Potts model, which was used successfully to model alignments of large collections of homologous sequences (Baldassi *et al*., 2014; Ekeberg *et al*., 2014, 2013; Morcos *et al*., 2011; Muntoni *et al*., 2021; Weigt *et al*., 2009). We apply this method to the natural viral diversity that accumulated over the first five years of SARS-CoV-2 circulation in humans.

## II. RESULTS

The basic rationale for estimating the fitness effects of mutations from their number of independent occurrences in the phylogenetic tree is that lineages carrying deleterious mutations will have fewer descendants that die out quickly, and are thus less likely to be sampled compared to lineages with neutral or even beneficial mutations (Bloom and Neher, 2023). The ratio of the number of occurrences of a mutation to its expected number under neutral evolution can be used as a quantitative proxy for the fitness effect (see Fig. 1B) (Haddox et al., 2025b) – see Sec. A.1 for details of the estimation procedure.

The SARS-CoV-2 phylogeny is characterized by distinct clades of very similar sequences separated by long branches along which the Spike protein in particular has undergone more dramatic changes. Motivated by the observations that diversity within clades is much smaller than across clades (Fig. 1C) we estimated fitness effects within each clade on a nearly constant background defined by the clade founder sequence. Differences between fitness effect estimates in different clades can then be due to (i) random fluctuations in observed counts; (ii) explicit temporal variation of selective pressure, and consequently of the viral fitness landscape; (iii) underlying epistatic interactions between protein residues. Explicit temporal variation in the fitness landscape is expected to be driven primarily by adaptation of the host immune response (Meijers *et al*., 2023).

As selection is expected to mainly act at the protein level, we focus on fitness effects of amino acid mutations throughout the rest of this work, dealing with fitness landscapes of individual structural proteins and particularly Spike. An analysis of the nucleoprotein is included in the supplementary material (Fig. S9). Each protein is represented as a chain of residues *i ∈* {1, …, *L*}, where *L* denotes the protein length and *σ*_*i*_ *∈* {*A, C*, …, *Y* } the amino acid identity. Founder sequences are in addition allowed to include gaps (*σ*_*i*_ = −). Our analysis then focuses on fitness effects of amino acid substitutions 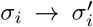 in a given clade *a*: 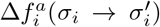. Fitness effects and their uncertainties are estimated using the computational pipeline available at (Haddox *et al*., 2025a). Our input data are the nucleotide mutation counts produced by the original pipeline (Bloom and Neher, 2023) applied to the UShER mutation-annotated tree from April 2024.

### A. Changing fitness effects

The left panel of Fig. 2A displays a scatter plot of estimated mutational effects in the Spike protein for two clades: 21J (Delta) and 21L (Omicron BA.2). Although most fitness effects are in good agreement with each other, there are some notable exceptions as for example the highlighted mutation at position 384. Substituting an L for the wild type amino acid P is beneficial in Delta and deleterious in 21L/BA.2. Such changes in fitness effects can arise from epistatic interactions between the position of interest and changes in the genetic background. Fig. 2B explores the fitness effects of P → L at position *i* = 384 across different clades and shows that the fitness effects cluster in two groups that align with the amino acid identity at position *j* = 408. If the amino acid at position *j* = 408 is a Serine, mutation P → L at position *i* = 384 is very deleterious, while if position *j* is an Arginine, the same mutation is neutral or slightly beneficial. Our goal here is to (i) systematically quantify the extent of changes in fitness effects, and (ii) explain these changes in terms of epistatic interactions with residues that differ between clades.

**FIG. 2.**
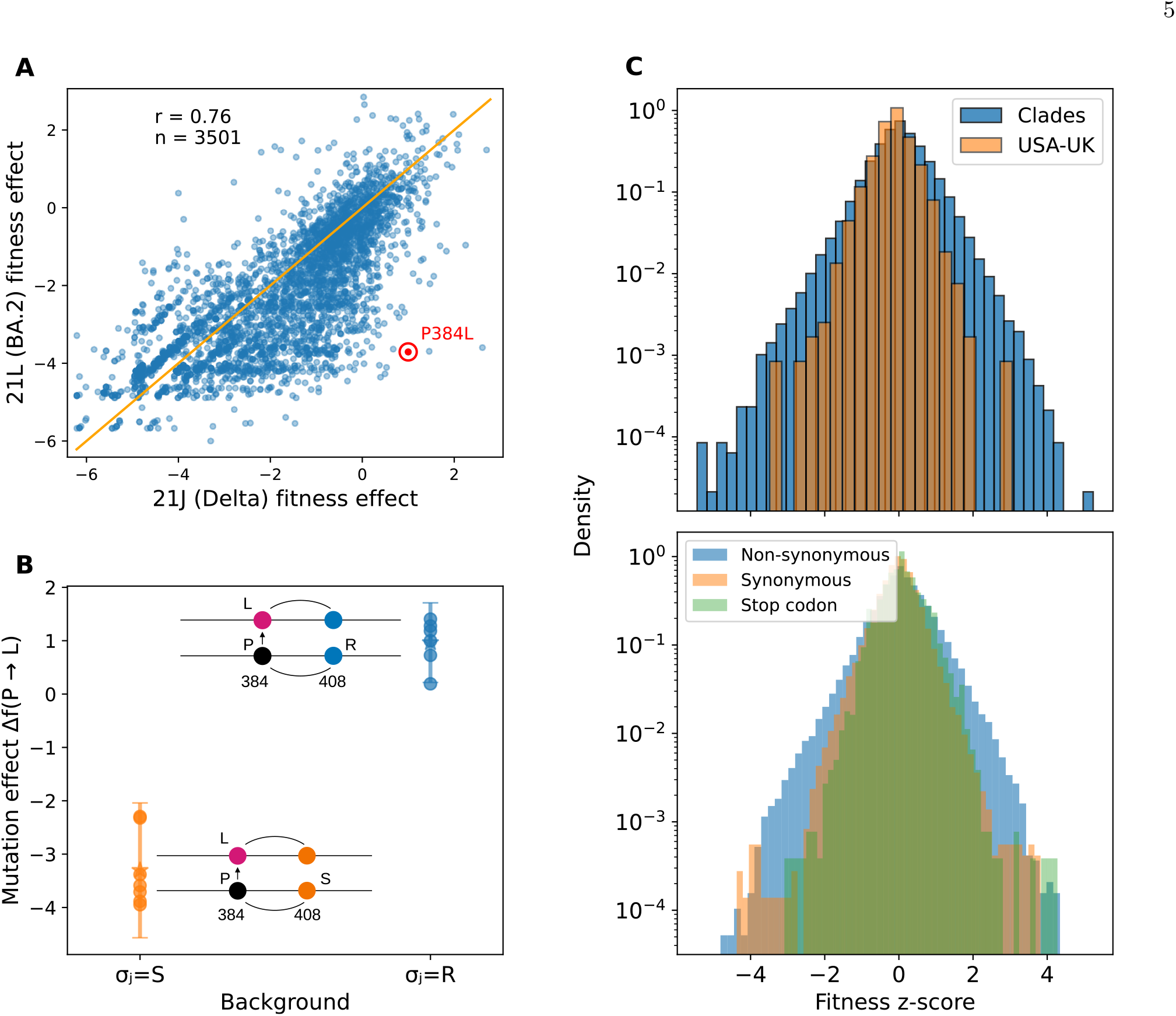
Fitness estimates and epistasis. **(A)** Scatter plot of fitness effects of mutations of the Spike protein between clades 21J (Delta) and 21L (Omicron BA.2). The highlighted red point corresponds to mutation (P384L) with strikingly different fitness effects in the two clades. **(B)** Fitness effects of mutation P *→* L at residue 384 in the Spike protein cluster in two groups separated by the amino acid identity at residue 408. A plausible explanation for the observed differences is the presence of epistatic interactions between the mutated residue at position 384 and changed background at position 408 as illustrated in the panel. The *⋆* symbol and the bar represent respectively the average and its standard deviation within the group. **(C)** Top, histogram of the mutational fitness z-scores for the Spike protein computed by: (blue) comparing the effect of a mutation in pairs of different clades; (orange) comparing the mutational effect as obtained from data belonging to different regions, i.e. USA and England. Bottom, histogram of z-scores (Eq. (1)) across clades over the whole set of structural proteins, stratified by: non-synonymous (blue), synonymous (orange) and early stop-codon (green).

To quantify the difference in the effect of a mutation in the genetic background of clades *a* and *b*, i.e. the epistatic signal, we compare mutational effects using a signed z-score:

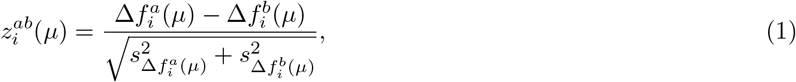

where 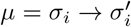 identifies an amino acid mutation, 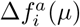 its fitness effect in clade *a*, and 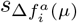 the associated uncertainty. Details about the definitions of 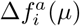 and 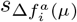can be found in Eq. (2A).

To further assess whether the observed differences in fitness effects are due to noise or epistasis, we compared the distribution of z-scores from Spike fitness effects estimated for the same clade using data only from the USA and England for clades 20I (Alpha), 21J (Delta), 21K (Omicron BA.1), 21L (Omicron BA.2), and 22B (Omicron BA.5). The USA and England have been mostly vaccinated with the same antigen and experienced similar waves of infection. We thus assume that differences in estimates within the same clade between the USA and England can be used as an estimate of the noise level. The distribution of z-scores for the comparison across clades is broader than that of the comparison of the same clade between the USA and England, see Fig. 2C (top panel). Notably, the tails of the distribution are much more pronounced for the comparison across clades. This inflation in the tails of the distribution of z-scores is likely due to changing fitness effects due to a combination of epistatic effects and temporal shifts in the fitness landscape.

The bottom panel of Fig. 2C shows the histogram of z-scores stratified over all structural proteins, i.e. Spike (S), Nucleocapsid (N), Membrane (M) and membrane Envelope (E), broken down according to the mutation type: non-synonymous 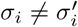 synonymous 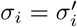 and nonsense 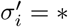 As expected, the distribution of z-scores for non-synonymous mutations has longer tails than that of synonymous mutations that don’t change the protein sequence.

Like-wise, stop-mutations that truncate the protein are expected to be equally deleterious across genetic backgrounds. There are some synonymous mutations with apparently high z-scores, but the majority of them are artefacts of variable sequence alignment in the vicinity of insertions or deletions. Taken together, these observations suggest that there is considerable epistatic modulation of fitness effects of amino acid mutations in different SARS-CoV-2 clades.

Next, we quantified shifts of fitness effects between two clades *a* and *b* by averaging absolute z-score at site *i* over all observed mutations at that position

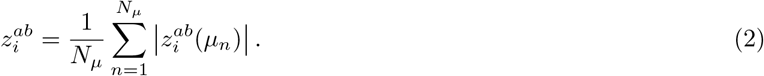

This site specific z-score for the Spike glycoprotein is shown in Fig. 3A for four pairs of clades. The horizontal axis spans the protein residues, while the vertical one displays the magnitude of the z-score. Visual inspection of the z-score traces along the protein suggests that the z-scores are elevated in the vicinity of mismatches between the clade pairs (indicated by vertical yellow bars). We quantified this observation using structures of the whole Spike trimer in different conformations (Bangaru *et al*., 2020; Benton *et al*., 2020; Ke *et al*., 2024; Shi *et al*., 2023; Zhang *et al*., 2021a,b). For those residues that were not included in the experimental structures, we used an AlphaFold (Jumper *et al*., 2021) model of the full trimer structure. We define the distance between two residues as the minimum distance between heavy atoms across the various structures and chains of the trimer. We then asked whether sites with elevated z-scores tend be closer to mismatching residues between the two clades compared to two different random expectations. Specifically, for each pair of clades, we computed the fraction of sites with 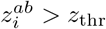 which are closer to a mismatching background residue than a threshold radius *r* = 5 or 10Å and plot this as a function of the threshold *z*_thr_. As expected, positions with high *z*-scores are more likely to be near mismatches. We also compared this with *random* benchmarks, obtained by redistributing the mismatching residues to random positions along the protein chain, or alternatively to all positions that have an alignment entropy above 0.01 in an alignment of around 4000 sequences samples approximately evenly in time and space. Sites with high 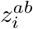 are more likely to be near mismatches in the background sequence than the random benchmark, especially for large z-scores and small radii (Fig. 3B).

**FIG. 3.**
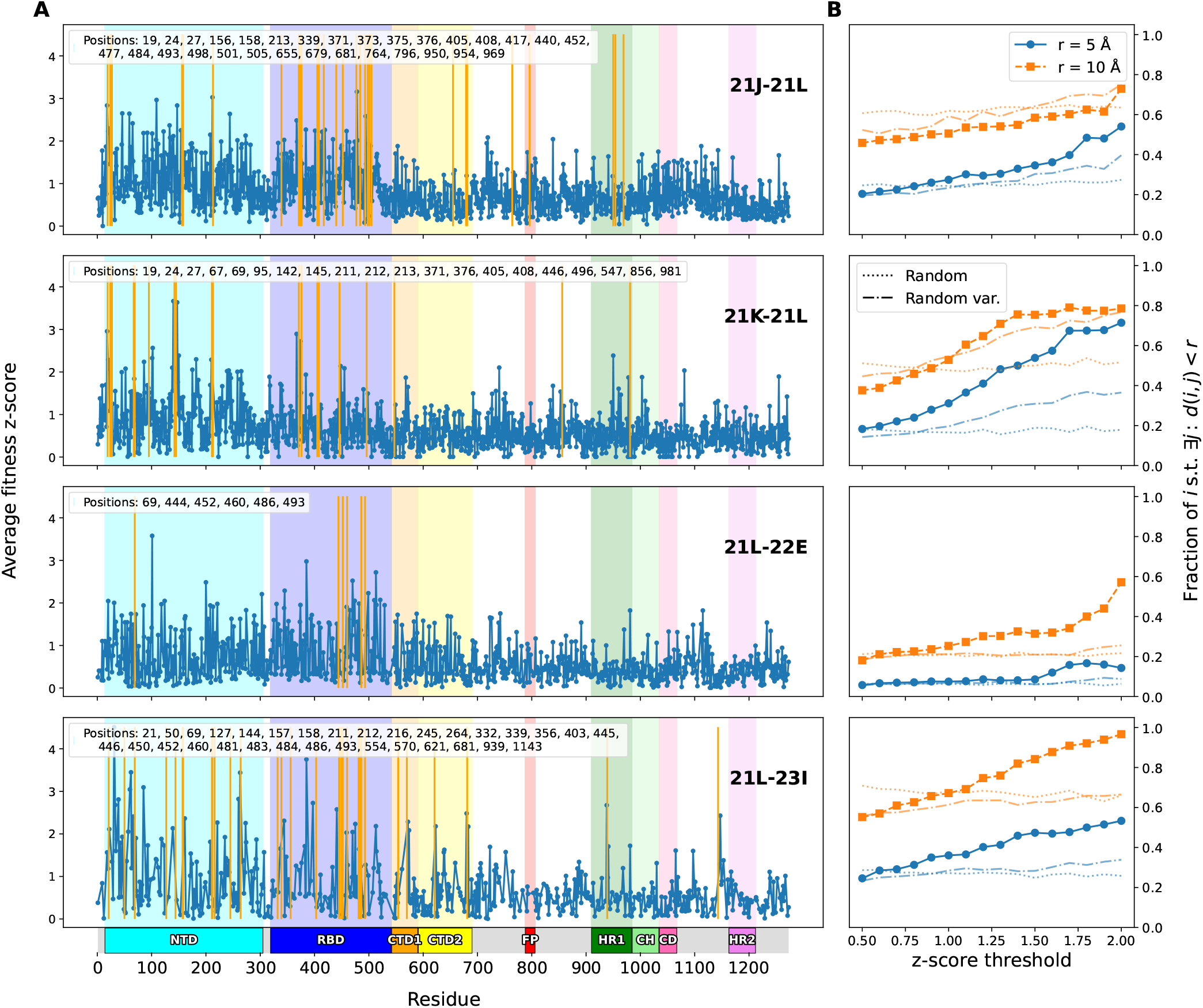
Spatial distribution of shifted fitness effects. **(A)** The plots shows z-scores (Eq. (2)) along the Spike protein sequence for four different clade pairs. From top to bottom: 21J-21L (Delta-Omicron BA.2), 21K-21L (Omicron BA.1-Omicron BA.2), 21L-22E (Omicron BA.2-Omicron BQ.1), 21L-23I (Omicron BA.2-Omicron BA.2.86). The solid yellow vertical lines highlight residues at which the founders of the two clades differ (with positions shown in the insert of each plot). Key functional domains of the Spike protein are highlighted and match the colors used in Fig. 5A. **(B)** Fraction of residues with z-score above threshold which also have a background mismatch within a sphere of radius *r*. The plots show the trend for increasing *z*-scores for each pair of clades for two choices of the sphere radius. The dashed and dotted lines in each plot show a random benchmark obtained by randomly drawing mismatches along the protein chain (dotted) or by randomly drawing from variable sites (dashed). By doing so, we aim to avoid picking random sites at which clade defining mutations are unlikely.

### B. Model

Epistatic interactions between residues have been successfully modelled using frameworks from statistical physics known as generalized Potts models (Morcos *et al*., 2011; Weigt *et al*., 2009). These models describe the probability of observing a particular sequence by a Boltzmann distribution parameterized by an “energy” which is proportional to the negative logarithm of the probability. This energy has terms for additive contributions of individual residues independent of the genetic background and pairwise epistatic interactions between residues. These models are typically inferred from large alignments of homologous sequences and have been used successfully to predict contact patterns between residues. The inferred models have also been used to predict fitness effects of amino acid substitutions in different genetic backgrounds (Figliuzzi *et al*., 2016) and we reason that this modeling framework is suitable to describe differences in fitness landscapes between different SARS-CoV-2 clades.

The fitness effect of a mutation 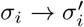 at site *i* can be expressed as its intrinsic (additive) effect *h*_*i*_ plus the sum of pairwise interactions *J*_*ij*_ with a set of residues *j* that are interacting with *i*:

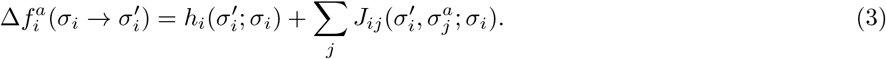

The interaction matrix *J*_*ij*_ will typically be sparse since residue *i* interacts only with a few partners *j*.

In Eq. (3), we explicitly indicate the dependency of the parameters on the starting amino acid *σ*_*i*_. For the rest of the manuscript, we will not specify this dependence explicitly for readability and simplicity. Note that the model is time independent and thus not able to capture explicit time variations of the mutational effects (see Sec. III for further limitations).

To estimate the model parameters, we compare all estimates of fitness effects in different clades. The differences in fitness effects between clades *a* and *b* for the same mutation 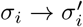 can be expressed as

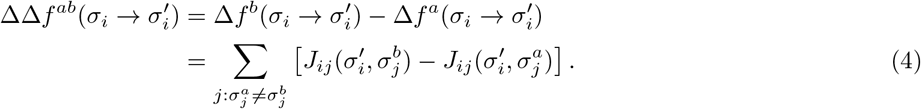

By taking the difference between pairs of clades we are able to remove the contribution of the additive effect *h*, which does not depend on the genetic background. In Eq (4), the summation runs over all sites *j* of the protein where the amino acid differs between clades *a* and *b*.

There are many more *J*_*ij*_(*σ*_*i*_, *σ*_*j*_) parameters than constraints from fitness estimates in different clades and inferring *J*_*ij*_(*σ*_*i*_, *σ*_*j*_) thus requires prior knowledge or some form of regularization. One reasonable assumption is that most residues do not interact, i.e. *J*_*ij*_(*σ*_*i*_, *σ*_*j*_) = 0 for most pairs (*i, j*), which can be achieved by minimizing the following objective function:

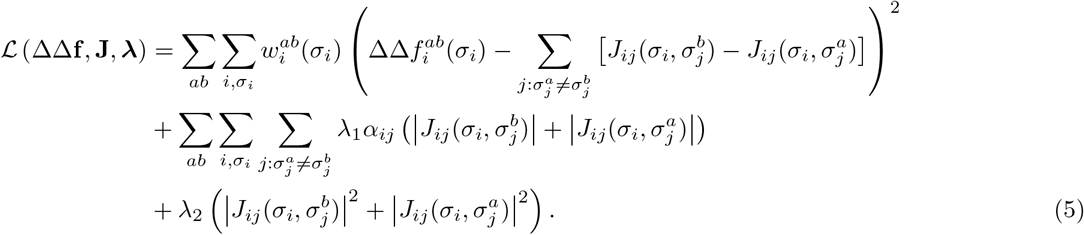

The first line is a loss *E*(**ΔΔf**, **J**) that enforces Eq. (4) in a soft way and can be interpreted as the mean squared error (MSE) of the model fit. The two outer sums therein run over the clade pairs and the observed mutations, and the weight 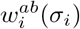of each term is inversely proportional to 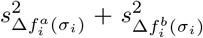, the combined uncertainty of 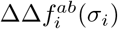 (we specify here only the dependence on the arrival amino acid *σ*_*i*_). The subsequent lines represent a cost (or regularization) contribution, and the hyperparameters *λ*_1_ and *λ*_2_ define the regularization strength. The overall resulting regularization is known as *elastic net* (Zou and Hastie, 2005), and it includes both *l*_1_ and *l*_2_-norm contributions. The *l*_1_-norm encourages sparsity of the model parameters. The additional multipliers *α*_*ij*_ allow to tune the *l*_1_ regularization strength differently for each residue pair (*i, j*). In particular, one can use a *biologically informed*prior, such that *α*_*ij*_ is proportional to the distance in three-dimensional space between residue *i* and *j*.

As long as *λ*_2_ *>* 0, the objective function is strictly convex, and the unique minimum can be efficiently found using gradient descent numerical techniques, see Methods. The optimal set of interaction parameters **J**^*\**^ are then given by **J**^*\**^(**ΔΔf**, ***λ***) = argmin_**J**_*ℒ* (**ΔΔf**, **J, *λ***).

### C. Model inference

While optimization of the convex objective function in Eq. (5) is straightforward, the hyperparameter *λ*_1_ needs to be chosen such that the *J*_*ij*_ capture epistatic shifts without overfitting noise. A clear signal of fitting noise would be a loss that is smaller than what would obtained when comparing estimates from the USA and England for the same clades as in Sec. II.A and Fig. 2C. This baseline loss is computed as 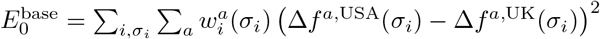,where the sum over *a* runs over the clades listed in Sec. II.A, and the weights 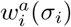 are defined as in Eq. (5).

Fig. 4B reports optimized losses *E*(**ΔΔf**, **J**^*\**^) for different choices of the *l*_1_ regularization strength *λ*_1_ and for the various functional forms of *α*_*ij*_ (Eq. (7A)), along with the baseline loss 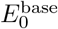 as a horizontal black line. As *λ*_1_ increases, more interactions are set to zero, causing the optimized loss to increase until 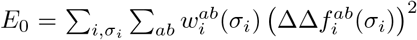 is reached when all **J** = 0. From this perspective, the *optimal* value of *λ*_1_ is the one that explains as much fitness variation as possible while still yielding 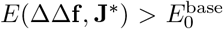, i.e., above the noise baseline. For the linear distance-based regularization *α*_*ij*_ = *d*_*ij*_*/d*_max_, this condition is met for *λ*_1_ *≳* 0.001. In particular, for *λ*_1_ = 0.001 the inferred interactions reduce the energy by 60% compared to *E*_0_ with only a 2.5% fraction of non-zero parameters, as shown in the bottom panel of Fig. 4B. Similarly, for the sigmoid-based *α*_*ij*_, the condition is met for *λ*_1_ *≳* 0.0002. These two regularization strength values are highlighted in Fig. 4B as vertical dotted lines. For further details about the inference algorithm and the choice of the parameters, we redirect the reader to Sec. A.2.

**FIG. 4.**
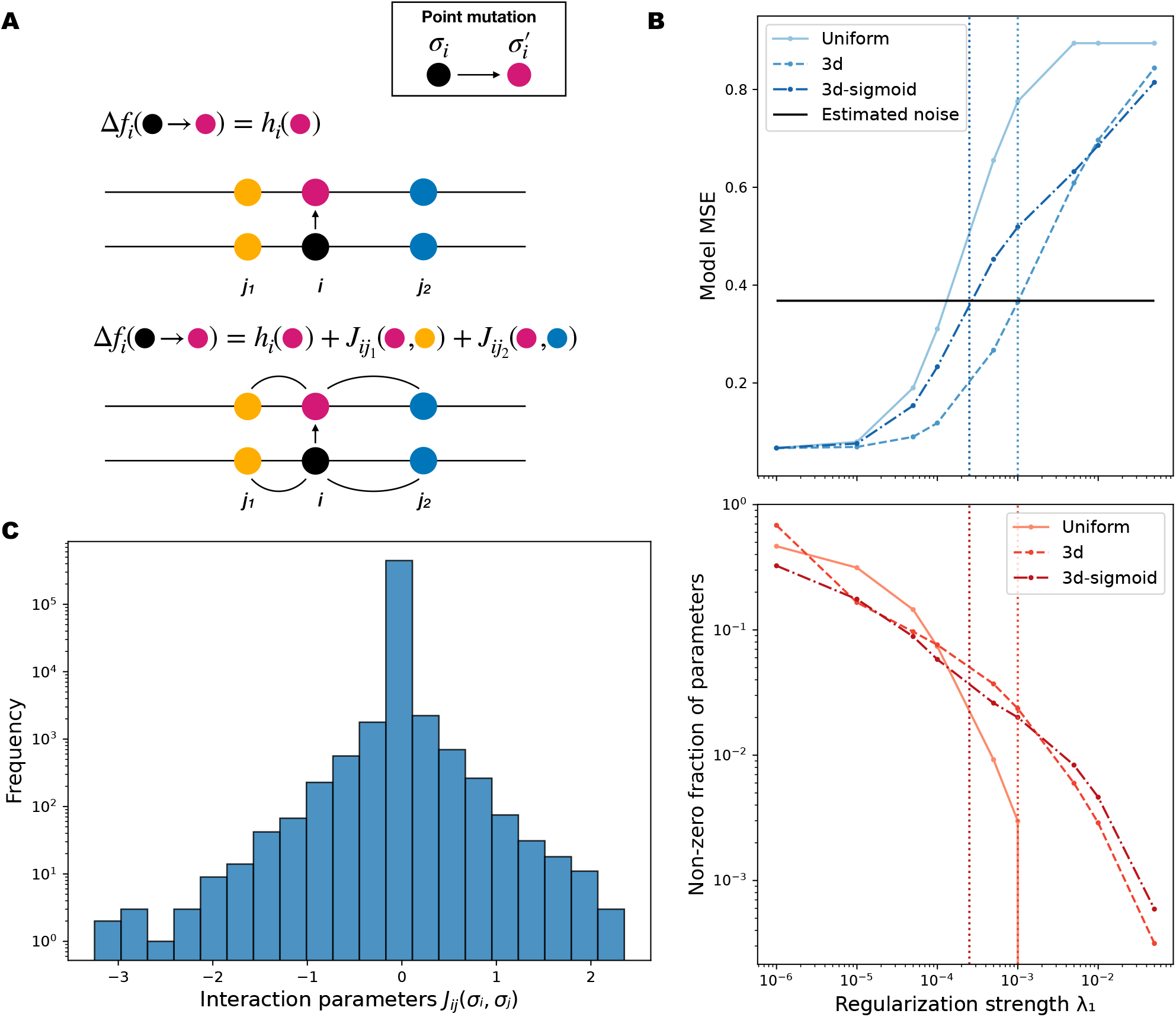
Model inference for the Spike protein. **(A)** Different models of mutational fitness: top, site independent model which does not account for interactions; bottom, pairwise epistatic model which makes the mutational effect context dependent. **(B)** Dependence of the model fit on the *l*_1_-norm regularization strength *λ*_1_ for different choices of the multipliers *α*_*ij*_ . Top, loss contribution to the objective function. The horizontal black line is a baseline estimate of the loss which can be ascribed to random fluctuations; bottom, fraction of non-zero interaction parameters. The two vertical dotted lines represent regularization strength values for which the distance-aware regularization strategies display mean squared errors right above the noise baseline estimate. **(C)** Log-scale histogram of the inferred interaction parameters for the 3d linear distance *α*_*ij*_ and *λ*_1_ = 0.001.

The uniform *l*_1_-norm regularization *α*_*ij*_ = 1 suffers from an inherent parameter degeneracy if certain background mutations are present in the same combinations in all clade pair comparisons. As a simple example, consider a change in fitness effect of a mutation *µ* for two pairs of clades (*a, b*) and (*c, d*). Both clade pairs differ at two residues *j*_1_ and *j*_2_ so that 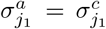 and 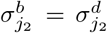 . The model can accommodate effect differences through the parameters 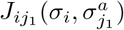 and 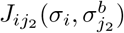 . However, without further information, the method is not able to discriminate which between residues *j*_1_ and *j*_2_ is responsible for the variation of fitness in the two contexts, and will thus assign the same absolute value to both interactions 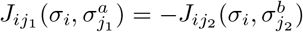. In SARS-CoV-2 evolution, it is common for variants to share many background mutations and this degeneracy is widespread. It can be broken by a distance dependent *α*_*ij*_, which is motivated by the observation and prior expectation that changing fitness effects tend to localize near mismatching background residues.

### D. Analysis and visualization of the inferred epistatic parameters

For our primary analysis, we use the linear distance-based regularization with *λ*_1_ = 0.001 based on a tree of GISAID data from April 2024 jbloomlab/SARS2-mut-fitness. The corresponding mutation counts and fitness effect estimates are available in the archived repository (Sesta and Neher, 2026).

More than 97% of the inferred epistatic parameters are zero and only a small fraction have large magnitudes, i.e. |*J*_*ij*_(*σ*_*i*_, *σ*_*j*_)| *≳* 1, which aligns with the biological expectation (see Fig. 4C). Fig. 5A shows an interaction map between protein residues based on the inferred epistatic interactions where the strength of residue-wise interaction score is computed as root-mean-square of parameters between all amino acids and positions 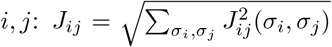 Note that the map is not symmetric: interaction parameters quantify the change of the fitness effect of mutations at site *i* for different contexts at site *j*. The reverse is often not observed, that is site *i* never changes between different clades. The inferred interaction parameters recapitulate the structural organization of functional domains of the Spike protein. Note, however, that this domain structure is enhanced by the distance-based regularization. The spatial clustering of inferred interactions is particularly evident for the N-terminal domain (NTD) and the receptor binding domain (RBD), which are characterized by an elevated density of background-defining mutations. There are also a few notable examples of inter-domain long-range interactions. These pairs are usually in spatial proximity across different chains of the Spike trimer.

**FIG. 5.**
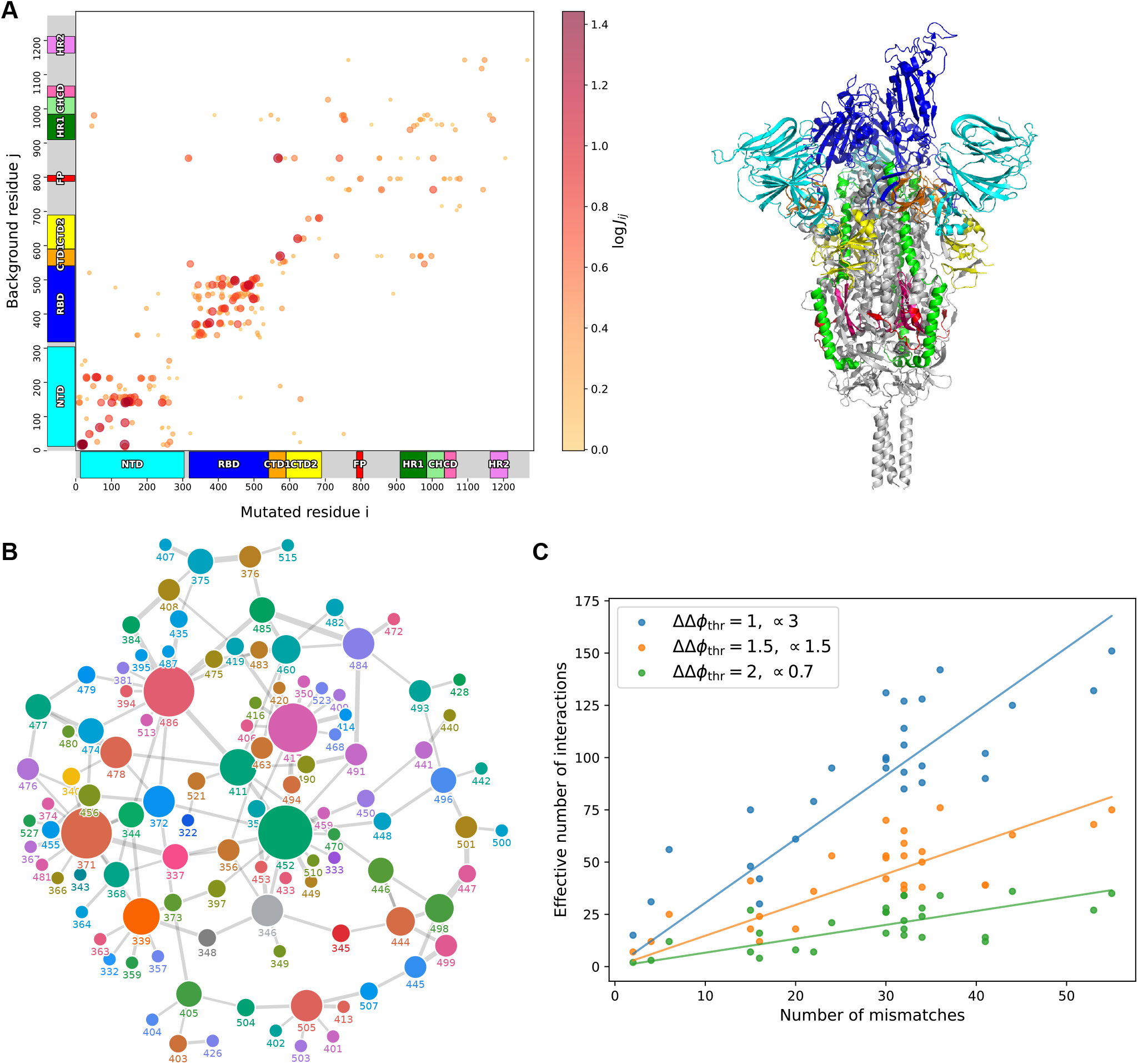
Visualization of inferred epistatic interactions. **(A)** Interaction map between protein residues. Each dot represents a specific residue pair (*i, j*). The color and size indicate the logarithm of the root-mean-square of the interactions *J*_*ij*_ . Only the 400 residue pairs with the highest *J*_*ij*_ are shown. Colored boxes highlight functional domains along the protein residues on both axes, while the plot insert shows the structural organization of the Spike trimer, with protein domains highlighted following the same color scheme. The folded structure is obtained from the PDB 7krr (Zhang *et al*., 2021b), representing the so called 1RBD-up conformation. **(B)** Interaction network between protein residues of the RBD, with interaction strength *J*_*ij*_ *>* 1.25. Circles represent residues, and their size is proportional to the number of links they share. Moreover, the thickness of the links is proportional to the strength of the interaction. **(C)** The plot shows the effective number of interacting partners as a function of the distance between clade pairs, computed as the number of mutations *µ* such that 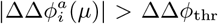, for three different choices of the threshold which are reported in the legend, together with an estimate of the slope of a fitted linear curve. The latter provides an estimate of the number of interactions per mismatch. The clade pairs are chosen from the following list of viral clades: 20I, 21J, 21K, 21L, 22B, 22E, 23A, 23I.

Fig. 5B shows a network representation of the inferred interactions of residues belonging to the RBD. In the network, sites 371, 417, 452, 486 have the largest number of epistatic interactions. Position 371 differentiates pre-Omicron from Omicron clades, where clade 21K (BA.1) was characterized by S371L while BA.2 and all subsequent Omicron descendants carried the substitution S371F. Position 417 was mutated in the early variants of concern Beta and Gamma as well as all Omicron variants. L452R is a defining mutation of Delta and the Omicron clades 22A (BA.4), 22B (BA.5) and 22E (BQ.1), while 452W fixed in the latest clades. Site 486 displays one of the highest diversity among clade founders, with four main different amino acids: F (wildtype), V, S and P. Sites 417, 452, 486 are directly involved in ACE2 binding (Lan *et al*., 2020; Liu et al., 2024) and immune evasion (Greaney *et al*., 2022). Site 371 interacts strongly with 455 through an inter-chain contact, where the latter is also involved in binding with ACE2 and immune escape. Among others, residue 452 interacts with 346, which is also known to be a site of strong immune escape. The strong interactions between groups of residues 444-448 and 496-501 inferred by our analysis have also been shown experimentally (Moulana *et al*., 2022; Starr et al., 2022a,b). The former are sites of strong immune escape, particularly for non-recombinant pre-BA.2.86 Omicron clades. The latter have been shown to have a strong impact on ACE2 binding, specifically mutations Q498R and N501Y (Starr *et al*., 2022a).

The interaction network of sites in the NTD is reported in Fig. S5. We identify residues 19, 69, 142, 145, 156, 158, 213 and 216 as the main centers of interactions. Mutational effects in the NTD have not been experimentally characterized as well as in the RBD. However, some studies have investigated immunogenic properties of mutations (Cerutti *et al*., 2021; Klinakis et al., 2021; Suryadevara et al., 2021), as well as their influence on binding with ACE2 (Dadonaite *et al*., 2025; Kugathasan et al., 2023), specifically through long-range interactions with the RBD in the full trimer configuration of the Spike. In particular, some protruding loops that are exposed on the surface of the protein collectively form the so called antigenic *supersite*, which is the main target of the host immune response focused on the NTD. The majority of the aforementioned residues either directly belong to the supersite, or are found on the edges of the involved loops. It appears that deletions at various positions of the supersite are well tolerated, as likely promoting immune escape, and defined major clades. For instance, deletions Δ69 − 70 and Δ144 first appeared in clades Alpha and 21K (BA.1) and showed very dynamic behavior since. Our analysis suggests that deletions at these sites have also a considerable epistatic impact, thus reshaping the virus mutational landscape.

In addition to inferring specific epistatic interactions, we sought to quantify the perturbation a typical change in the genetic background causes and asked the following question: for each mismatch that alters the genetic background, how many mutational effects are affected? We considered a subset of clade pairs corresponding to major clades and the associated set of shared mutational effects. For each clade pair, we computed the number of mutations for which the quantity 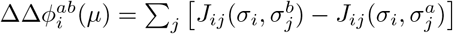 exceeds a chosen threshold in absolute value, and plot this number as a function of the distance between the clades (Fig. 5C). The resulting trend is approximately linear, consistent with the hypothesis that each additional mismatch introduces roughly the same number of interacting partners. This number can be estimated as the slope of a linear fit constrained to pass through the origin. To assess the sensitivity of this estimate to the threshold choice, we repeated the analysis using three different threshold values. We find that the inferred number of interacting partners decreases with increasing threshold, with slope values ranging from 3 to 0.7. This suggests that each additional mismatch typically induces appreciable shifts in the fitness landscape at around 1-3 positions.

### E. Validation of the inferred epistatic interactions

Fig 6A displays the relation between the inferred epistatic interactions and the discrepancies of fitness estimates for selected pairs of positions. Each panel shows the fitness effect of an amino acid mutation 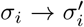 in different clades on the vertical axis and the interaction effects 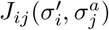 at the most strongly coupled site *j* on the horizontal axis. As expected, the inferred interactions are correlated with the average fitness effect within the groups of clades for all mutations reported in Fig. 6A. The panel corresponding to RBD mutation P384L, also shown in Fig. 2B, illustrates how the method was able to successfully explain the background dependent fitness via interaction with site 408 in state *σ*_*j*_ = S, R. The panels corresponding to mutations R21G, P499L and R683Q show that the model works well for cases in which the background can assume three possible configurations. For all three, the average fitness effects align well with the inferred interactions. The correlation between fitness effects across clades and interaction parameters holds when most of the modulation of the fitness effect can be a explained by a single interacting partner *j*, as is the case for 3 our of the 4 displayed scatter plots. However, if the mutation is inferred to have many interacting partners the relationship between fitness effects and interaction parameters to a specific partner is less clear. Mutation P499L (top right in Fig 6A) is a notable example. Site 499 is inferred to be interacting with several other residues besides 445, such as 444, 478 and 452. These additional interactions explain the large variation of P499L on background 445V, e.g. the much less deleterious effect in clades 21I, 21J and 22E. Examples of other mutations can be found in Fig. S6.

**FIG. 6.**
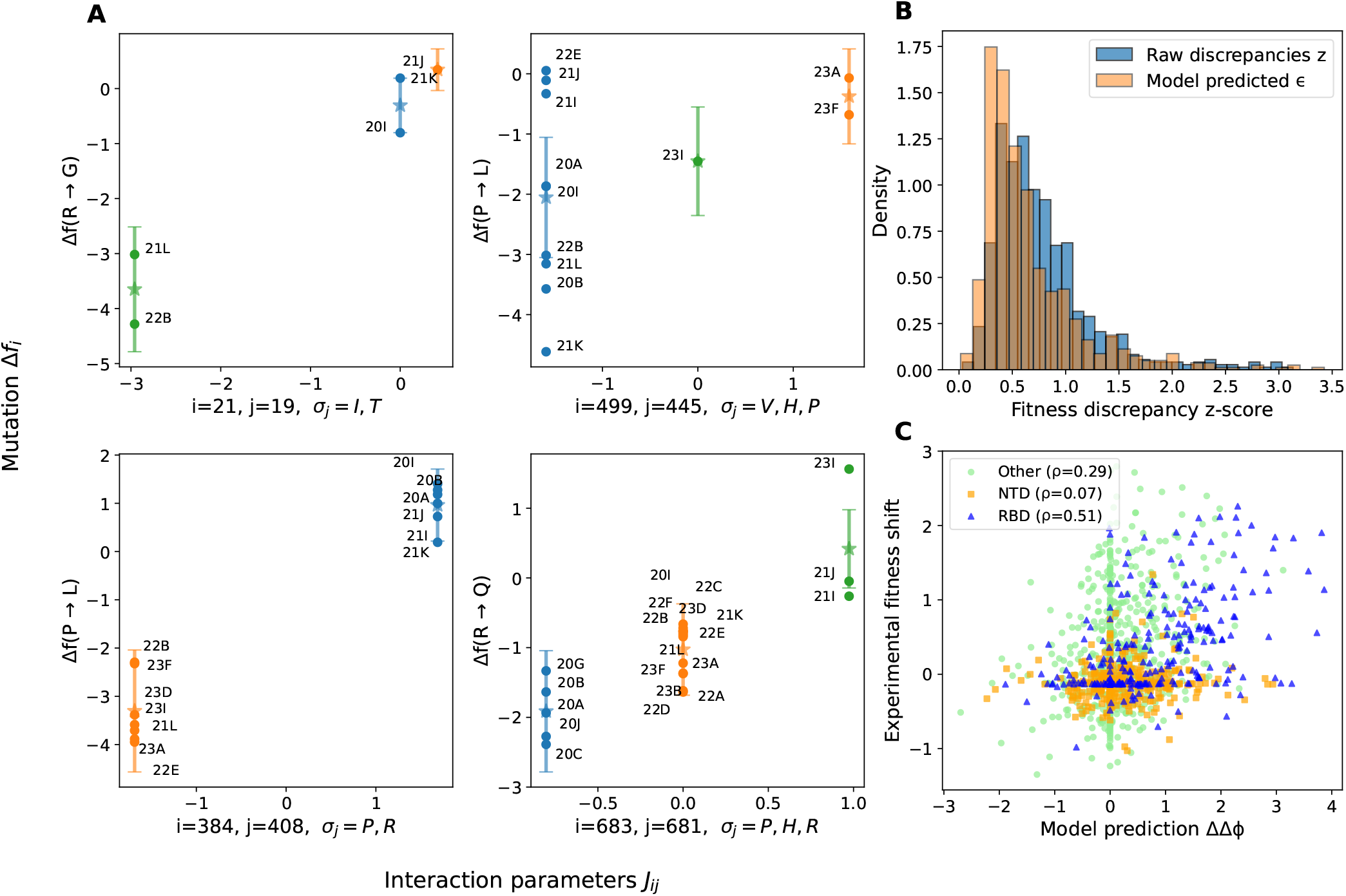
Validation of the inferred epistatic interactions. **(A)** The plots scatter the values of the inferred interaction parameters on the x-axis against fitness effects of mutations in different clades on the y-axis for strongly interacting residue pairs. The state variable background amino acid 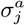 is indicated on the x-axis in order of increasing interaction strength. For each group of clades, the *⋆* symbol corresponds to the mean effect within the group, whereas the error bar indicates the standard error of the mean. **(B)** Histograms of raw and model predicted z-scores averaged over the available clade pairs (Eq. (7)), for the 705 mutations observed in the train and test clades. **(C)** Scatter plot between model predicted fitness discrepancies ΔΔ*ϕ* and corresponding estimates obtained from experimental measurements of the phenotype *cellular infection*. The clade pair chosen for the comparison is 21J-21K (Delta-Omicron BA.1), and the legend reports the Pearson correlation coefficient between the two quantities within different domains.

As a further validation, we split the Spike protein fitness data into a training and a test set. The training set includes clades from 20A to 23H (HK.3), while the test set consists of clades 23I (Omicron BA.2.86) and 24A (Omicron JN.1). Counts for clade 24A are taken from the public UShER tree as of November 2024. Clade 24A descended from 23I, which in turn emerged from 21L (Omicron BA.2) via a long branch (see top rightmost part of the tree in Fig. 1A – orange samples) (Khan *et al*., 2023). Clade 23I differs from 21L by 32 mismatches in Spike, including 20 novel mutations and 12 which are either reversions to the root or mutations observed elsewhere in the tree. Clade 24A, which eventually replaced 23I, carries an additional L455S mutation. Given this high divergence we expect multiple epistatic interactions, as illustrated in Fig. 3 for the clade pair 21L-23I. Our goal is to test whether interactions inferred from the training set can account, at least partially, for the changing fitness effects between the test clades and the training set.

To quantify the model’s predictive power, we compare the raw normalized fitness discrepancies 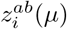 (see Eq. (1)) with the model-predicted ones:

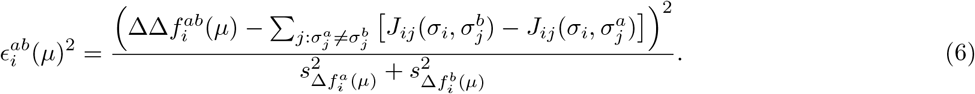

The quantity 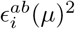 can be interpreted as a mutation and clade-pair-specific component of the loss (up to a normalization factor). Both raw and model-predicted z-scores can be averaged over clade pairs to obtain a mutation-specific score:

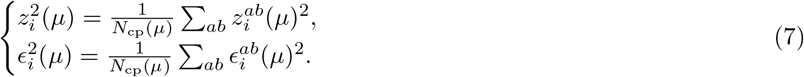

where *N*_cp_ is the number of available clade pairs for mutation *µ*. The analysis includes 705 mutations present in both training and test sets. For *∼*75% of them, the model reduces the raw z-score, i.e., *ϵ*_*i*_(*µ*) < *z*_*i*_(*µ*). This indicates partial predictive power despite the novel background mismatches in clades 23I and 24A not being present in the training set. Fig. 6B shows the histograms of raw and model-predicted discrepancies. The model-predicted z-scores are shifted toward lower values, with a median of 0.50 compared to 0.67 for the raw scores, highlighting a systematic reduction. Tab. I reports pairs of raw and model predicted scores 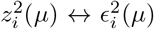 for the 10 mutations displaying the largest drop. Despite the improvement, in many cases *ϵ*^2^ remains significantly above zero, reflecting noise and interactions with background mismatches between training and test clades that are not represented in the training set alone. When the novel interaction is weaker in absolute value than the one inferred from known backgrounds, the model still provides a benefit. However, if the novel interactions dominate the model is uninformative.

**TABLE I.** Raw *z*^2^ and model predicted *ϵ*^2^ scores averaged over the available clade pairs, for the ten mutations with the largest z-score reduction *z*^2^ *− ϵ*^2^.

| $z_i^2(\mu)$ | $\epsilon_i^2(\mu)$ | Mutation |
| --- | --- | --- |
| 7.06 | 1.27 | T385I |
| 8.29 | 2.94 | L18F |
| 6.36 | 1.10 | P85L |
| 5.16 | 0.40 | G482S |
| 4.18 | 0.27 | T1027I |
| 3.92 | 0.03 | P384L |
| 7.68 | 4.10 | T20I |
| 8.99 | 5.60 | R683Q |
| 5.68 | 2.42 | R683W |
| 3.42 | 0.28 | P85S |

Finally, we compare the inferred interactions with independent estimates obtained from in-vitro DMS experiments. Dadonaite et al. (2023) have probed the impact of mutations across the entire Spike protein in several major viral variants on different phenotypes (Dadonaite *et al*., 2024, 2023, 2025). Mutational effects on the phenotype *cellular infection* – a combination of binding with ACE2 and cell entry – are available for variants 21J (Delta), 21K (BA.1) and 21L (BA.2). For variants 21L (BA.2) and 23A (XBB.1.5) ACE2 binding effects and cell entry effects are reported separately (Dadonaite *et al*., 2024). The more recent results of variant KP.3.1.1 (Dadonaite *et al*., 2025) can not be reliably compared to our estimates due the limited amount of sequence data available.

Bloom and Neher (2023) showed that DMS measurements of mutational effects on infectivity correlate well with fitness effects estimated from mutation counts in global sequence data with an overall Pearson correlation of 0.66. The unexplained variance is partly due to experimental noise, inference noise, and differences between the experimental phenotype and the in-vivo viral fitness. With measurements available for multiple variants, we can now test whether shifts of mutational effects in DMS experiments are also captured by our inferred model.

Before comparing experimental fitness discrepancies with our model predictions, we preprocessed the data through the `multidms` pipeline introduced in (Haddox *et al*., 2023). The method infers fitness mutational effects jointly combining measurements realized on different variants, at the same time determining shifts with respect to a chosen reference. For our purposes, we chose 21K (BA.1) as the reference for data from Dadonaite et al. (2023) and 21L (BA.2) for data by Dadonaite et al. (2024). We compare shifts in DMS data with shifts in fitness effects predicted by our model 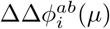 (see definition related to Fig. 5C). In Fig. 6C, we show the result of the comparison for the clade pair 21J-21K (Dadonaite *et al*., 2023), whereas scatter plots for clade pairs 21L-21K and 23A-21L (Dadonaite *et al*., 2024) can be found in the SI (Fig. S8). The model is once again inferred from the GISAID tree as of April 2024 (clades 20A to 23I). The shifts predicted by the model and experimental shifts are moderately correlated with each other, with Pearson coefficient 0.32. The experimental and estimated mutational fitness effects themselves are correlated with each other with correlations coefficients around 0.5 to 0.7 (see Figs. S3, S4). It is thus not surprising that correlation between model-predicted and experimental shifts between clade pairs is lower than the correlation between the experimental and estimated mutational fitness effects. In particular, regularization drives many interaction/shift parameters to zero, which results in a large number of points on one or the other axis in the scatter plot. However, we see that the model predictions are able to capture general trends. For instance, the Delta variant seems to be more tolerant to Spike mutations than BA.1, resulting in the predominance of points in the upper right quadrant in Fig. 6C. The majority of sites displaying large effects belong to the RBD, which we expect to be subjected to similar selective pressure both in the wild and in-vitro. Experimental shifts and model predictions in the RBD are also well correlated (*ρ* = 0.51), whereas the correlation is much weaker in the NTD (*ρ* = 0.07) and moderate elsewhere (*ρ* = 0.29).

## III. DISCUSSION

Previous work had shown how the landscape of fitness effects of single mutations away from a wild type genome can be accurately estimated from mutations observed in millions of sequenced viral genomes (Bloom and Neher, 2023; Haddox *et al*., 2025b). Here, we show that not only single mutation effects, but also interactions between mutations can be inferred from such data. This inference sheds light on how the fitness landscapes of SARS-CoV-2 proteins change while the virus evolves from year to year and accumulates mutations. Due to the limited diversity of SARS-CoV-2, this inference is restricted to interactions with sites that have changed between major viral clades, but nonetheless provides clear evidence of the epistatic interactions that shape the viral fitness landscape. While the idea is applicable to all viral proteins, most SARS-CoV-2 proteins have undergone limited recent evolution and there is thus limited potential for epistatic interactions. We therefore focused on the Spike protein of the virus which differs substantially between major clades.

Epistasis and the resulting shifts of mutation effects can give rise to diverging evolutionary trajectories and adaptive potential of different variants. The emergence of Omicron involved a major reconfiguration of key parts of the Spike protein (Moulana *et al*., 2022) and led to an era of rapid accumulation of mutations therein (Roemer *et al*., 2023). Limited epistasis implies that measurements such as deep mutational scanning (DMS) of a single variant can be used to predict the fitness effects of mutations in future variants. This question had been previously investigated by Doud et al. (2015) for NP proteins of H1N1 and H3N2 lineages of influenza. The authors found that mutational effects are mostly conserved between these two homologous proteins. A similar approach applied to the rapidly evolving envelope protein of HIV-1 revealed stronger evidence of shifting landscapes of mutational effects (Haddox *et al*., 2018). On much longer evolutionary time scales, epistasis and the resulting loss of predictability of evolution was investigated using DMS and reconstructed ancestral steroid receptors (Park *et al*., 2022). By comparing landscapes of fitness costs estimated from mutation counts in large collections of SARS-CoV-2 genomes across major variants that differ by 20-50 substitutions in the Spike protein, we find similar patterns of gradually shifting landscapes. As expected, and in line with previous work (Haddox *et al*., 2018), shifted effects tend to be in proximity of positions where the background variants differ from each other but can be long ranged as well. When fitting a pairwise interaction model, every substitution on average affects the fitness effect of 1.5 positions through an interaction term with magnitude greater than 1.5 (*∼* 5 fold difference in mutation count). This picture is qualitatively consistent with results by (Park *et al*., 2022) who found that sites with the highest rate of epistitic shifts randomize their mutational effects after sequence divergence of less than 50%.

Inferring pairwise interaction matrices from mutation count data is related to the problem of inferring Potts models from multiple sequence alignments of protein families (Morcos *et al*., 2011; Weigt *et al*., 2009). Instead of parameterizing a putative equilibrium distribution of sequences, our model parameterizes the rate at which mutations are observed in sequence data with a typical sampling density (Bloom and Neher, 2023). The model attempts to explain the observed fitness effects of mutations in different clades through pairwise interactions with background residues. The large number of parameters and the limited number of combinations in which sets of changing background residues are observed results in degeneracies and the necessity to regularize the inference. We break these degeneracies by using a distance-based regularization that encourages a sparse set of interactions with residues in spatial proximity to the mutated site. This prior encourages interactions to be between nearby residues, but this local enrichment of interactions is also supported by randomization tests (see Fig. 3).

Another intrinsic limitation of our approach is that the model assumes that the viral fitness landscape is time-invariant, whereas in reality it changes due to interactions with the host immune system. In such cases, the model may introduce spurious interactions to explain time-dependent fitness effects. Two strategies may help identify such cases: (i) examining the distribution of fitness effects across clades with identical backgrounds, where unexplained outliers may signal explicit temporal variation; and (ii) inspecting long-range interactions between residues that are far apart, which are more likely to be artifacts. Nevertheless, distant residues could still interact for example because they affect protein dynamics or stability. As a possible future extension, the model could be generalized to incorporate time dependence, for instance by allowing the additive contribution in Eq. (3) to depend on the clade index, 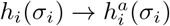. Such a time dependence could capture the changing immunity profile of the human population and the resulting shifting fitness landscape of the virus (Haddox *et al*., 2026; Meijers et al., 2023).

Inference of epistatic interactions from mutation counts complements DMS experiments on multiple backgrounds (Taylor and Starr, 2023, 2024) in that it depends on viral fitness across the entire transmission cycle. As sequencing of viral genomes becomes more widespread, data volumes will increase for many other viruses beyond SARS-CoV-2 and inference of fitness landscapes from sequence and mutation data will become increasingly feasible. For viruses with well defined subtypes or clades, the methods introduced here can help to understand how fitness landscapes shift as the virus evolves.

## ACKNOWLEDGMENTS

The fitness estimates used in this work depend on the pandemic scale phylogenetic trees maintained by the Angie Hinrichs in the UShER team at UCSC. This work builds on the open data sharing by 1000s of scientists and health care professionals around the world who have contributed SARS-CoV-2 sequences to NCBI or GISAID. We gratefully acknowledge stimulating discussion with Georg Angehrn and Cornelius Roemer. This work was supported by the Swiss National Science Foundation (SNSF) through grant 310030 188547.

## Appendix A Methods

### Data Availability Statement

All data used in this study are available in the repository associated with this manuscript https://github.com/neherlab/SC2Epistasis/, which is archived on Zenodo at https://doi.org/10.5281/zenodo.21107313. Genetic diversity data were obtained from the Nextstrain SARS-CoV-2 staging tree, by downloading the *Genetic Diversity Data (TSV)* file on May 22nd 2026. Additional external datasets are linked directly to their original sources.

#### 1. Bayesian fitness estimates

Given a nucleotide mutation *x* → *y* with *x, y ∈ N* = {*A, T, C, G* } at site *k* of the genome, the fitness effect of the mutation is estimated from the following quantities:

- 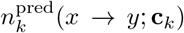: the predicted number of counts from the neutral evolution model, which depends on the specific site *k* and its condition **c**_*k*_ (Haddox *et al*., 2025b).
- 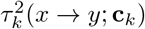:residual variation unexplained by the neutral evolution model, which represents the uncertainty on the predicted counts.
- 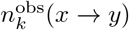 the number of independent mutation counts actually observed on the phylogenetic tree.

These quantities are subsequently fed into a simple Bayesian framework, which provides the posterior probability distribution for the mutational fitness effect. From there, the fitness estimate and uncertainty are given by:

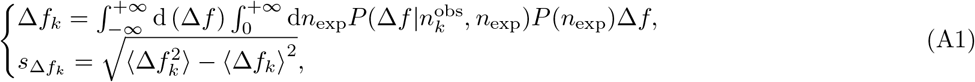

where we dropped the mutation dependency for readability and both Δ*f* and *n*_exp_ are dummy variables to be integrated over. Specifically *n*_exp_ is a log-normally distributed random variable with mean log 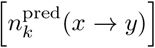 and variance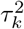.

The fitness score obtained so far refers to nucleotide mutations. To obtain fitness effects of amino acid mutations, estimates of nucleotide substitutions producing the same amino acid substitution are aggregated through a weighted average:

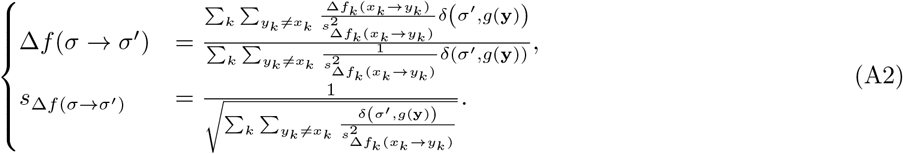

In Eq. (2A) the sum over *k* runs over the nucleotide positions belonging to the codon which encodes for the residue of interest, whereas the sum over *y*_*k*_ runs over all nucleotide such that *y*_*k*_ ≠ *x*_*k*_. The function *g*(**y**) maps the codon to the corresponding amino acid *σ*^*′*^, so that the delta function retains only the *y*_*k*_’s producing the desired amino acid mutation. Furthermore, Δ*f*_*k*_(*x*_*k*_ → *y*_*k*_) and 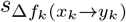 are defined according to Eq. (1A).

#### 2. Inference algorithm

For a collection of clade pairs for which the effect of mutation 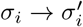 is observed, the set of equations as in Eq. (4) can be rewritten in a compact matrix form:

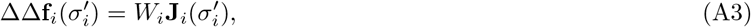

where the matrix *W*_*i*_ has elements:

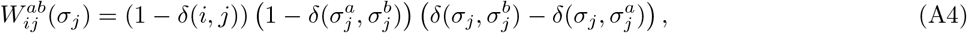

with *δ*(*i, j*) and *δ*(*σ*_*i*_, *σ*_*j*_) being discrete Kroenecker delta functions. Without the *l*_1_-norm regularization the problem is solvable analytically since the objective function becomes a quadratic form of the interaction parameters. For a single mutation, this reads:

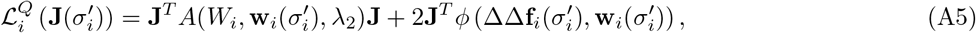

where *A* 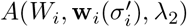 and 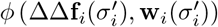are the quadratic form matrix and shift vector respectively. As long as *λ*_2_ *>* 0 the matrix *A* is positive definite, and thus the problem can be solved by:

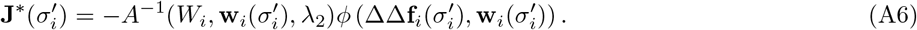

However, when *λ*_1_ ≠ 0 one has to resort on numerical methods to optimize the objective function in Eq. (5). Specifically, we used an orthant-wise limited memory quasi-Newton (OWL-QN) gradient descent algorithm, which is specialized for objective functions that include *l*_1_-norm regularization. Indeed, for large values of the *l*_1_ regularization strength *λ*_1_, the objective function becomes non-differentiable for *J ≃* 0, which is the case for most components of the interaction parameters. We implemented the algorithm in Julia, by generalizing the code that can be found at https://gist.github.com/yegortk/ce18975200e7dffd1759125972cd54f4.

In Sec. II.C, we display the outcome for three choices of *α*_*ij*_:

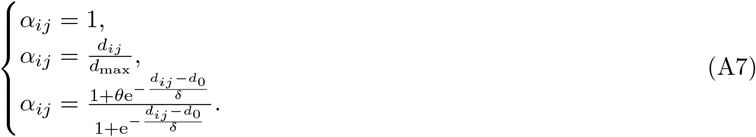

The first choice corresponds to a uniform regularization strength. The second one rescales *λ*_1_ proportionally to the distance between *i* and *j*, while the third is a sigmoid function of the distance and depends on a few extra hyperparameters. Specifically, *d*_0_ represents the typical range of interaction among protein residues, which is around 10Å, while *δ* is the decay length scale that defines how sharp the shape of the sigmoid function is. The results shown in Fig. 3 help rationalizing the choice of regularization multipliers *α*_*ij*_ which encourage sparsity based on the distance between protein residues. In order to implement the two distance dependent regularization strategies, i.e. second and third line of Eq. (7A), we used several PDB structures of the full Spike protein in different conformations and calculated the minimal distance between heavy atoms of pairs of residues. Tab. II reports the complete list of PDB’s. In addition to this, we also used a model structure obtained via AlphaFold to cover those residues that are not included in any of the PDB’s.

**TABLE II.**
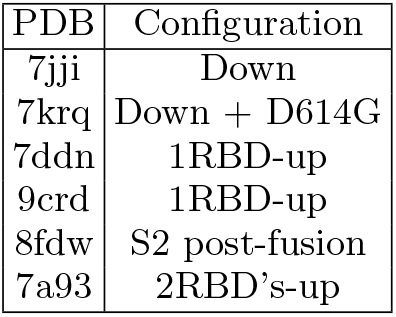
List of PDB’s used to compute distances between protein residues in the Spike trimer.

The linear distance regularization strategy only depends on *d*_max_, the maximum distance between residues across all PDB’s, which amounts to *d*_max_ = 217Å. On the other hand, the sigmoid-like *α*_*ij*_ depends on the following hyperparameters: *θ, d*_0_, *δ*. To perform the inference, we chose such hyperparameters as: *θ*=0.01, *d*_0_=15Å, *δ*=1.25. To gain some intuition on this regularization strategy, let’s consider the asymptotic limits as a function of the distance *d*_*ij*_. For *d*_*ij*_ *∼* 0 one has *α*_*ij*_ *∼ θ*, and so the overall regularization scales as *λ*_1_*θ*. On the other hand, when *d*_*ij*_ *≫ d*_0_, *α*_*ij*_ → 1 and thus the regularization converges to *λ*_1_.

As stated in the main text, the *l*_2_ regularization guarantees that the objective function in Eq. (5) is strictly convex, and thus a unique solution exists. The parameters initialization we chose is **J**=0.

#### 3. The multidms pipeline

The `multidms` method combines available experimental replicates and mutational effects measured on the backgrounds of different variants in order to infer a global epistatic model. The *shift* parameters in the model quantify epistasis between mutations. These are subject to a sparsity-encouraging *l*_1_-norm regularization. For the detailed documentation, see https://matsen.group/multidms/index.html. By jointly modeling mutational effects in different conditions, `multidms` allows to infer more reliable estimates, partially removing the effect of random fluctuations and condition biases among different experiments.

To compare our model predictions to mutational fitness effects measured in Dadonaite et al. (2023), we directly accessed the table at https://github.com/matsengrp/SARS-CoV-2_spike_multidms/blob/main/results/spike_analysis/mutations_df.csv, which reports functional scores for the three variants: 21K (reference), 21J and 21L.

To obtain refined values of experimental mutational effects from Dadonaite et al. (2024), we downloaded raw variant-wise functional scores from https://github.com/dms-vep/SARS-CoV-2_Omicron_BA.2_spike_ACE2_binding/blob/main/results/func_scores/Lib1-221023_high_ACE2_func_scores.csv (21L) and https://github.com/dms-vep/SARS-CoV-2_XBB.1.5_spike_DMS/tree/main/results/func_scores (23A). Then, we inferred the global epistatic model through the `multidms` pipeline choosing *λ* = 10^−5^ for the *l*_1_-norm regularization strength.

