## Supplementary Information for "Epistasis and the changing fitness landscapes of SARS-CoV-2"

### Supplementary Information for: Detecting epistasis from SARS-CoV-2 genome data

Luca Sesta
Richard A. Neher

August 17, 2026

#### Figures and Tables

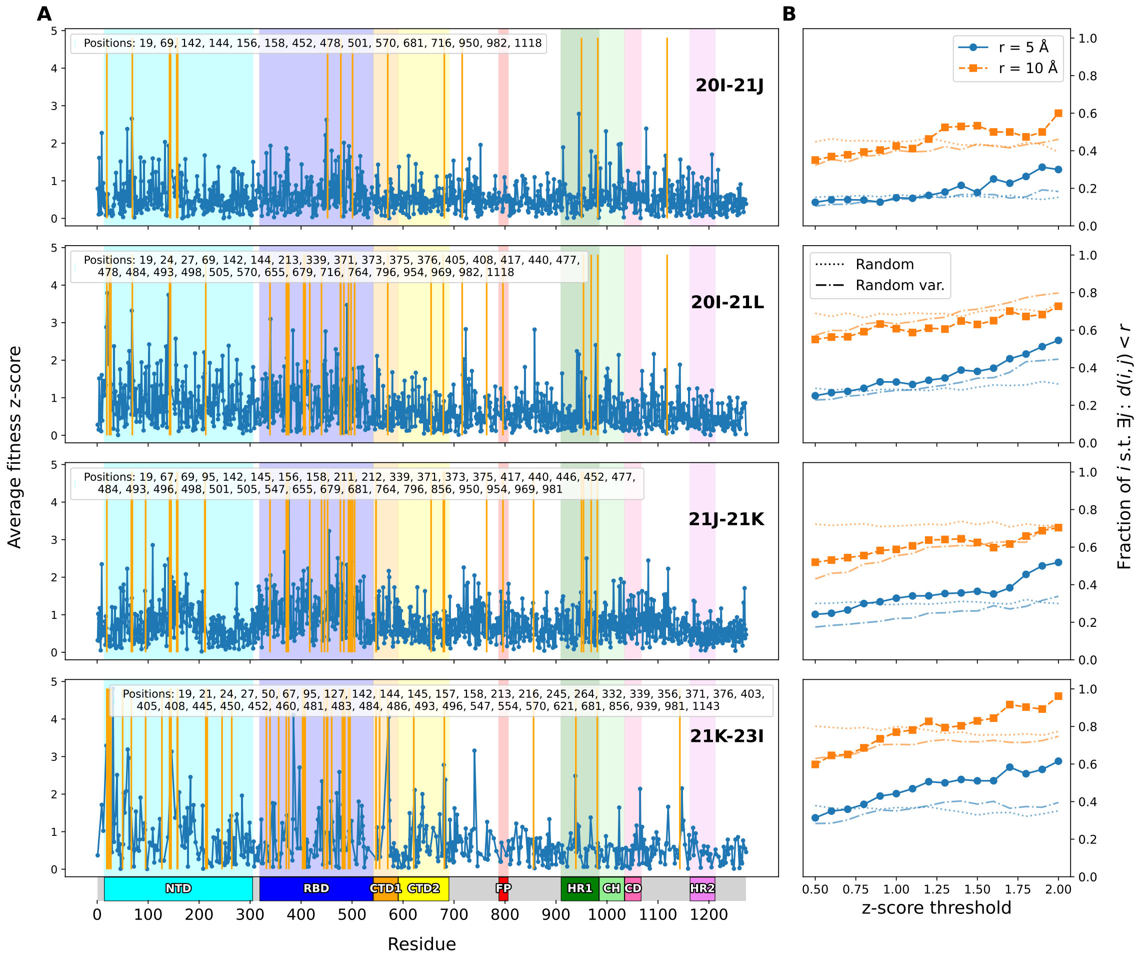


**Figure S1: Raw epistatic signal for clade pairs: 20I-21J (Alpha-Delta), 20I-21L (Alpha-Omicron BA.2), 21J-21K (Delta-Omicron BA.1), 21K-23I (Omicron BA.1-Omicron BA.2.86). (A)** Scatter plot of site specific z-scores (Eq. (2)) against residue indexes of the Spike protein. The solid yellow vertical lines highlight mismatching residues between each pair of clades, and the list is also reported in the insert of each plot. The colored boxes below the plot highlight some key functional domains of the Spike protein. **(B)** Fraction of residues with z-score above threshold which also have a background mismatch within a sphere of radius $r$. The plots show the trend for increasing $z$-scores for each pair of clades for two choices of the sphere radius. The dashed and dotted lines in each plot show a random benchmark obtained by randomly drawing mismatches along the protein chain (dotted) or by randomly drawing from variable sites (dashed). By doing so, we aim to avoid picking random sites at which clade defining mutations are unlikely.

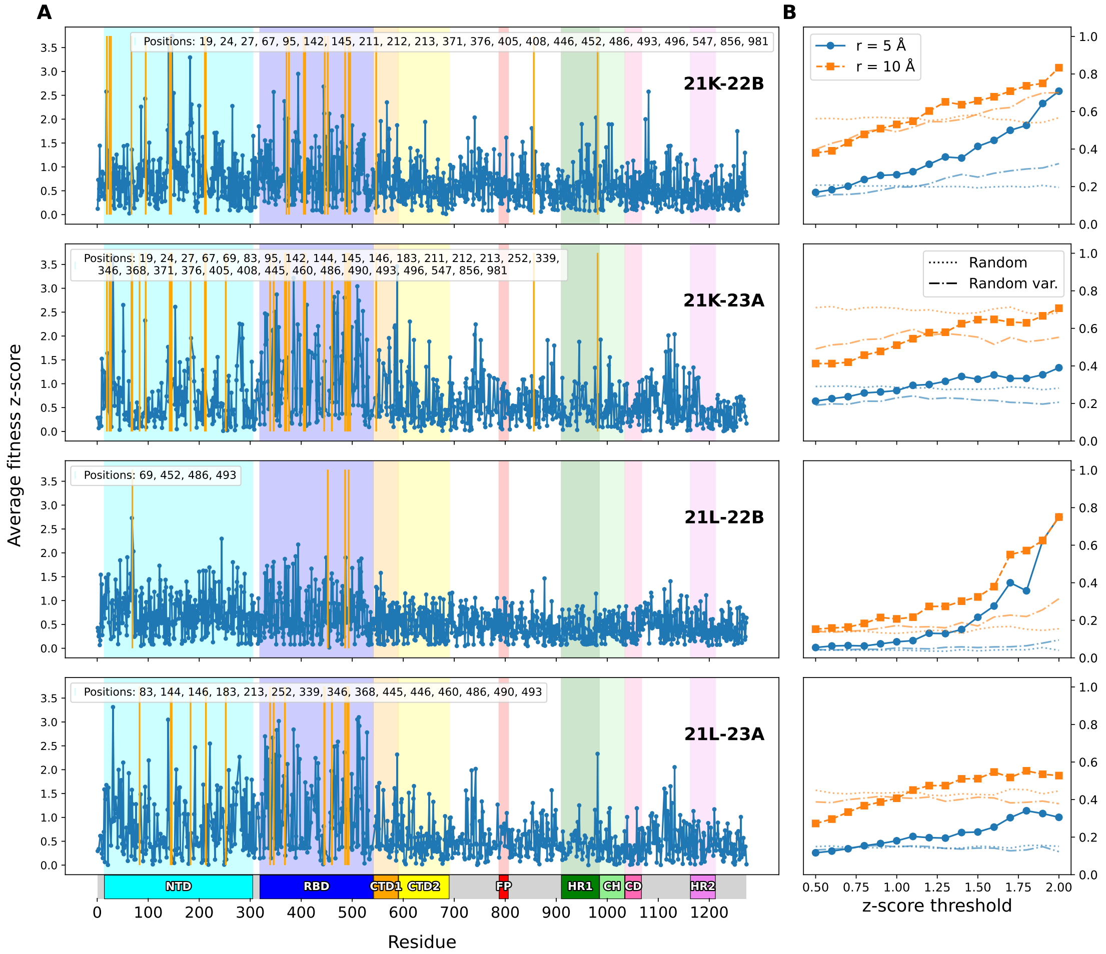


**Figure S2: Raw epistatic signal for clade pairs: 21K-22B (Omicron BA.1-Omicron BA.5), 21K-23A (Omicron BA.1-Omicron XBB.1.5), 21L-22B (Omicron BA.2-Omicron BA.5), 21L-23A (Omicron BA.2-Omicron XBB.1.5). (A)** Scatter plot of site specific z-scores (Eq. (2)) against residue indexes of the Spike protein. The solid yellow vertical lines highlight mismatching residues between each pair of clades, and the list is also reported in the insert of each plot. The colored boxes below the plot highlight some key functional domains of the Spike protein. **(B)** Fraction of residues with z-score above threshold which also have a background mismatch within a sphere of radius $r$. The plots show the trend for increasing $z$-scores for each pair of clades for two choices of the sphere radius. The dashed and dotted lines in each plot show a random benchmark obtained by randomly drawing mismatches along the protein chain (dotted) or by randomly drawing from variable sites (dashed). By doing so, we aim to avoid picking random sites at which clade defining mutations are unlikely.

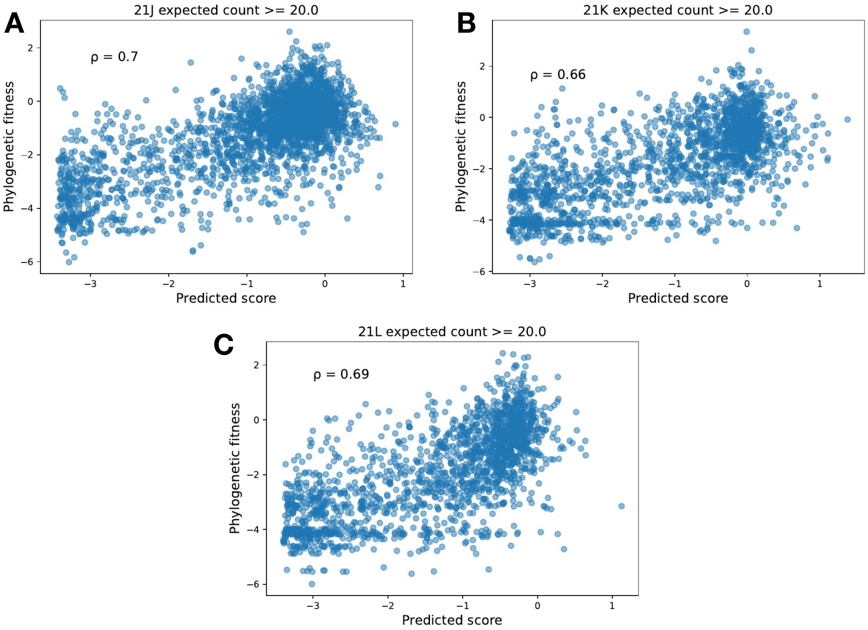


**Figure S3: Scatter plots between phylogenetic and predicted fitness.** Comparison between mutational fitness effects estimated from: phylogenetic data (vertical axis), DMS on pseudoviruses [1] (horizontal axis). Fitness effects from DMS data are jointly preprocessed through the `multidms` pipeline [2]. The strain chosen as reference is 21K. Pearson correlation coefficients are reported within the plots.

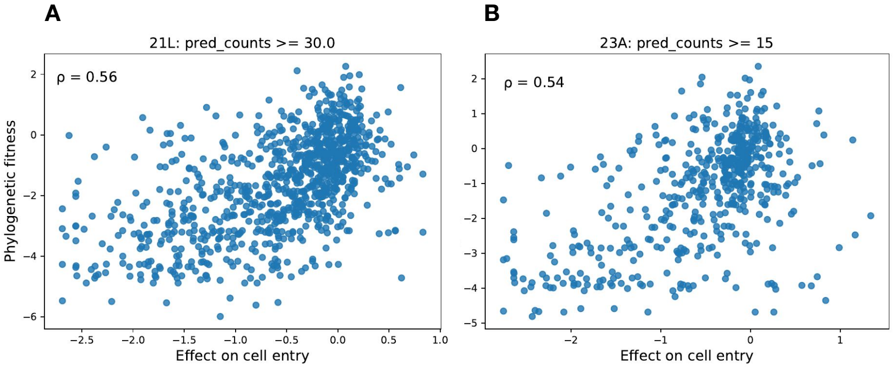


**Figure S4: Scatter plots between phylogenetic and experimental effect on cell entry.** Comparison between mutational fitness effects estimated from: phylogenetic data (vertical axis), DMS on pseudoviruses [3] (horizontal axis). Fitness effects from DMS data are jointly preprocessed through the `multidms` pipeline [2]. The strain chose as reference is 21L. Pearson correlation coefficients are reported within the plots.

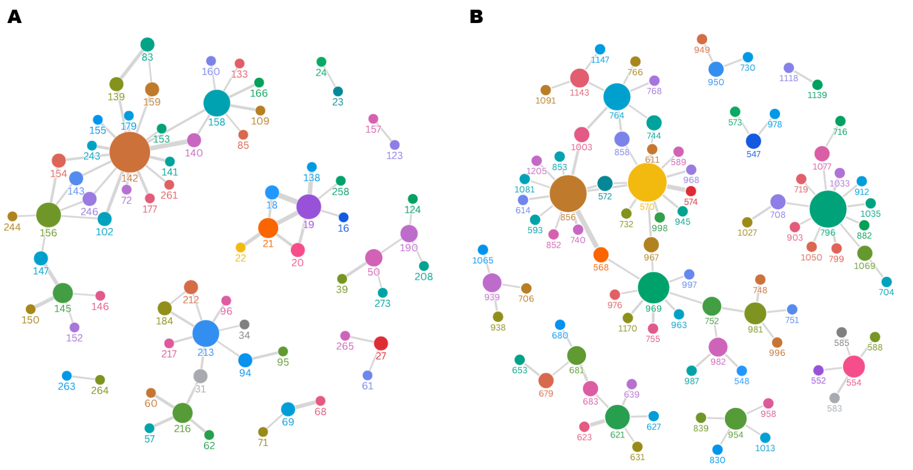


**Figure S5: Residue-residue interaction networks for NTD and S2 + S1/S2 junction.** The size of the circles is proportional to the number of connections going through it, whereas the width of a link is related to the coupling strength. **(A)** Interaction network for the NTD, retaining residues for which $J_{ij} > 1.25$. **(B)** Interaction network for the S2 and S1/S2 juction regions, retaining residues for which $J_{ij} > 1.25$.

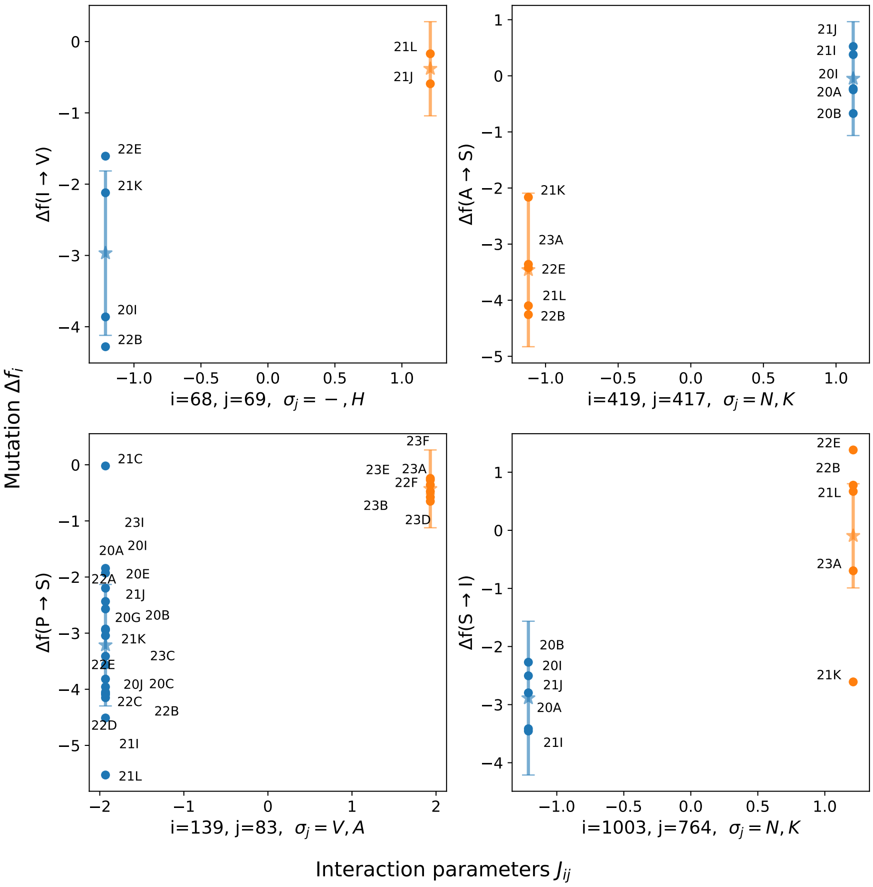


**Figure S6: Inferred couplings align with changing fitness effects.** The plots scatter the values of the inferred coupling parameters against mutational fitness effects of clades carrying the corresponding background amino acid, for strongly interacting residue pairs. On the horizontal axis, the graphs show the values of the couplings $J_{ij}(\sigma_i', \sigma_j^a)$, with $\sigma_j^a$ varying for the different groups of clades, as reported on the horizontal axes labels. On the vertical axis, the graphs show the fitness effects of mutation $\sigma_i \to \sigma_i'$ in the different clades. For each group of clades, the $\star$ symbol corresponds to the mean effect within the group, whereas the error bar quantifies the standard deviation of the mean as $\sqrt{\sum_a s^2_{\Delta f_i^a(\mu)}/N_c}$, with $N_c$ the number of clades in the group.

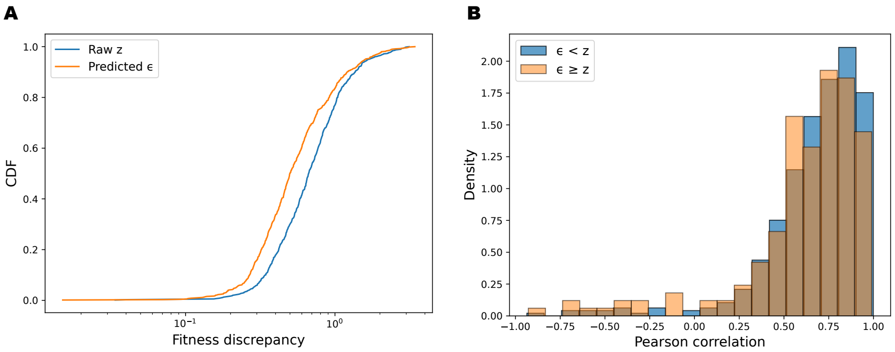


**Figure S7: Further results on couplings validation. (A)** Cumulative Distribution Functions (CDFs) for the raw $z$ and model predicted $\epsilon$. The offset between the two curves underlines a systematic reduction. **(B)** Histogram of the Pearson correlation coefficients between raw and model predicted mutation-wise z-scores, computed over the available clade pairs. The histogram is stratified between mutations for which injecting the model provides an improvement ($\epsilon < z$) and when it does not ($\epsilon \geq z$). The two distributions are strongly overlapping, underlying the significance of couplings inferred from previously observed background mutations and validating the additive nature of the pairwise epistatic model.

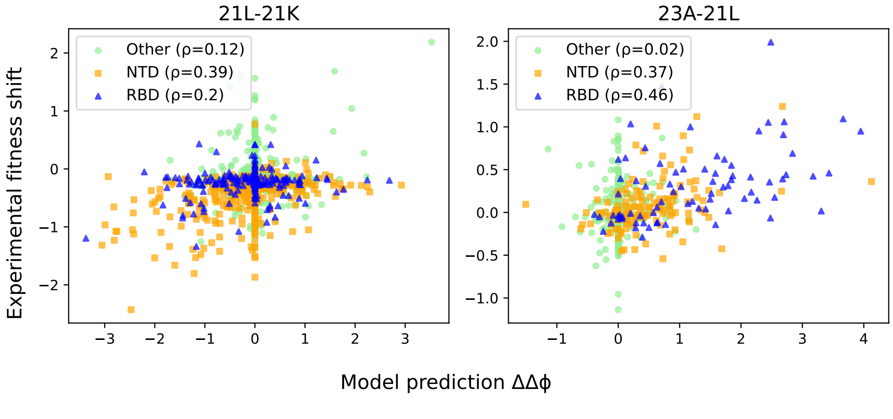


**Figure S8: Comparison between model predicted and independent experimental measurements of fitness discrepancies.** Left. Clade pair 21L-21K (Omicron BA.2 - Omicron BA.1). Right. Clade pair 23A-21L (Recombinant XBB.1.5 - Omicron BA.2). The legends report the Pearson correlation coefficients between the model predicted and experimental estimates.

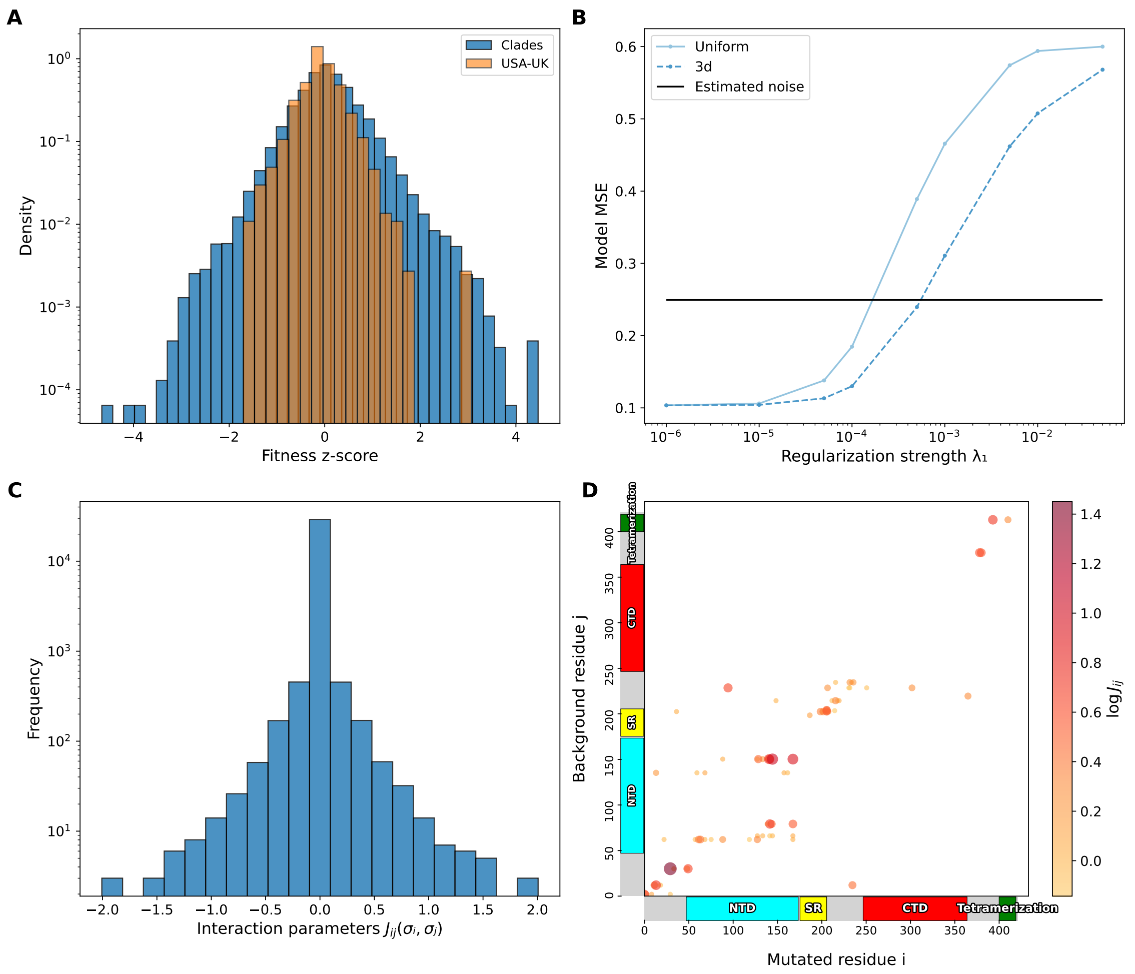


**Figure S9: Epistatis in the Nucleocapsid protein (A)** Histogram of the mutational fitness z-scores for the Nucleocapsid protein computed by: (blue) comparing the effect of a mutation in pairs of different clades; (orange) comparing the mutational effect as obtained from data belonging to different regions, i.e. USA and England. **(B)** Trend of the model mean squared error as a function of the regularization strength $\lambda_1$, for the uniform and 3d regularization strategies. The horizontal black line is computed from the total fitness discrepancies of the regional comparison, and stands as an estimate of noise. The distances used for the 3d regularization rely on the PDB structure of the monomer obtained from the AlphaFold model. **(C)** Histogram of the inferred coupling parameters with 3d regularization and $\lambda_1$=0.001. **(D)** Interaction map between protein residues. Each dot represents a specific residue pair $(i, j)$. The color indicates the logarithm of the Frobenius norm $J_{ij}$ of the associated couplings. Only the 75 residue pairs with the highest $J_{ij}$ are shown. Colored boxes highlight functional domains (or disordered domains like SR and Tetramerization) along the protein residues on both axes.

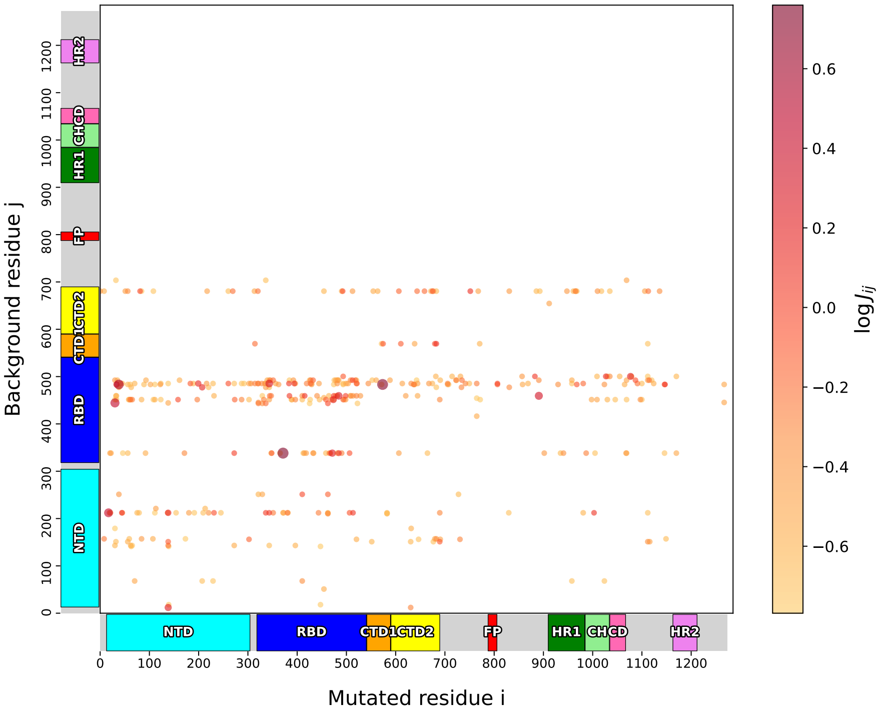


**Figure S10: Interaction map between protein residues inferred with uniform regularization.** The chosen $\lambda_1$ regularization strength is 0.00015, which corresponds to a model energy (mean squared error) slightly above the noise baseline. Each dot represents a specific residue pair $(i, j)$. The color indicates the logarithm of the Frobenius norm $J_{ij}$ of the associated couplings. Only the 400 residue pairs with the highest $J_{ij}$ are shown. Colored boxes highlight functional domains along the protein residues on both axes. Compared to Fig. 5A of the main text, it is apparent that the map showed here is strikingly affected by the coupling degeneracy related to irreducibly shared background mismatches. Given a mutation, we define the irreducible set of background mismatches as the ensemble of residues which assume the same two amino acid configurations across all clade pairs comparisons. Any putative fitness discrepancy signal that involves sites in the irreducible set is evenly spread over the whole set, and eventually concentrated on the few background residues which do not belong to it. This is why in the map we only observe a limited number of horizontal *stripes*, which correspond to those background mismatches that do not belong to irreducible sets.

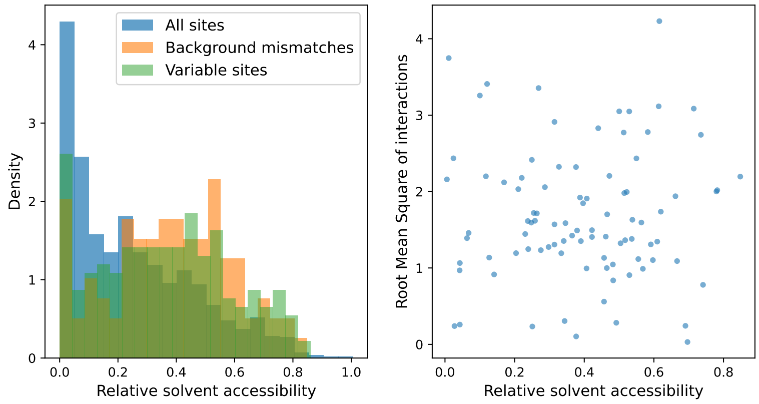


**Figure S11: Relative solvent accessibility (RSA) of variable and interacting residues.** **(A)** Distribution of RSA for: all residues (blue), positions which are mismatches between clade founder sequences (orange), variable positions with alignment entropy larger than 0.01 in a set of about 4000 sequences sampled evenly since the beginning of the pandemic (green). Variable positions are enriched for surface exposed residues. **(B)** Maximum of the root mean square of the interaction strength $J_{ij}$ over mutated sites $i$ for a given position $j$ that differ between clade founder sequences. Not clear relationship with RSA is observed.

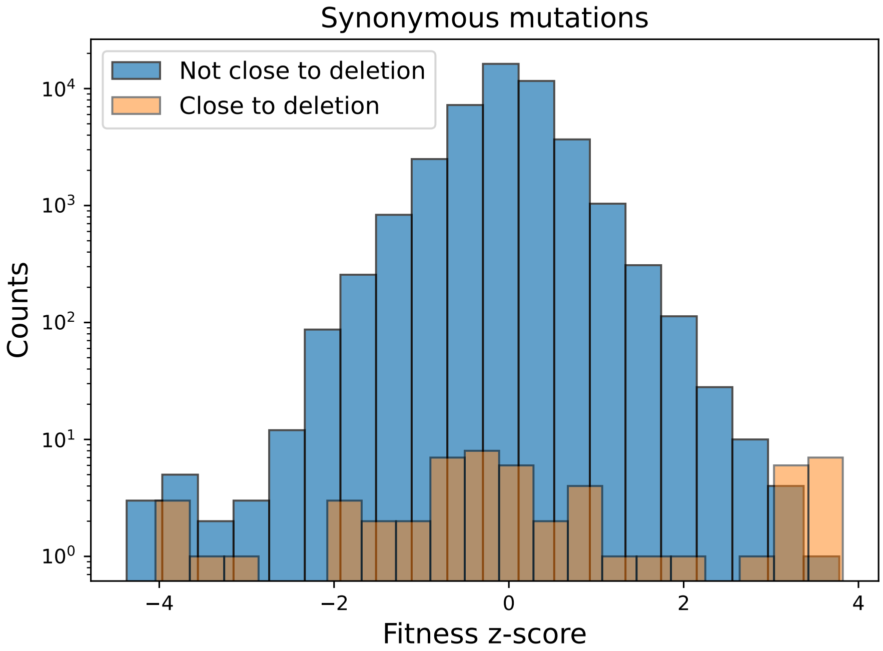


**Figure S12: Histograms of z-scores for synonymous mutations.** Blue: mutations for which none of the two clades in the comparison have a deletion in the vicinity of the mutated residue. Orange: mutations for which one or both clades in the comparison possess a deletion in the vicinity of the mutated residue. The proximity threshold is set to 6 nucleotides.
